# Longshot: accurate variant calling in diploid genomes using single-molecule long read sequencing

**DOI:** 10.1101/564443

**Authors:** Peter Edge, Vikas Bansal

## Abstract

Short-read sequencing technologies such as Illumina enable the accurate detection of single nucleotide variants (SNVs) and short insertion/deletion variants in human genomes but are unable to provide information about haplotypes and variants in repetitive regions of the genome. Single-molecule sequencing technologies such as Pacific Biosciences and Oxford Nanopore generate long reads (≥ 10 kb in length) that can potentially address these limitations of short reads. However, the high error rate of SMS reads makes it challenging to detect small-scale variants in diploid genomes. We introduce a variant calling method, Longshot, that leverages the haplotype information present in SMS reads to enable the accurate detection and phasing of single nucleotide variants in diploid genomes. Using whole-genome Pacific Biosciences data for multiple human individuals, we demonstrate that Longshot achieves very high accuracy for SNV detection (precision ≥0.992 and recall ≥0.96) that is significantly better than existing variant calling methods. Longshot can also call SNVs with good accuracy using whole-genome Oxford Nanopore data. Finally, we demonstrate that it enables the discovery of variants in duplicated regions of the genome that cannot be mapped using short reads. Longshot is freely available at https://github.com/pjedge/longshot.

## Introduction

The availability of second-generation DNA sequencing technologies such as Illumina short reads has made the resequencing of human genomes routine (1). Both SNVs, the most abundant form of variation in the human genome, and small indel variants can be reliably detected using whole-genome Illumina sequencing using sequence coverage of 30–40× (2; 3). Nevertheless, sequencing human genomes using short-read sequencing technologies has many limitations. First, humans are diploid organisms with two copies (maternal and paternal) of each autosomal chromosome. Haplotypes, or the sequence of alleles that occur on an individual chromosome, can be computationally assembled from whole-genome sequencing using overlaps between reads that span multiple heterozygous variants (4; 5; 6). However, due to the low rate of heterozygosity of human genomes (7), Illumina reads derived from paired-end sequencing of short fragment libraries (200–500 bp in length) typically cover only a single variant site, and do not provide long-range haplotype information. Second, approximately 3.6% of the genome consists of long and highly similar duplicated sequences where short-reads cannot be uniquely mapped and hence SNVs cannot be detected. These regions overlap hundreds of coding genes, including many disease associated genes such as PMS2 and STRC (8).

Third-generation single molecule sequencing (SMS) technologies such as Pacific Biosciences (PacBio) and Oxford Nanopore (ONT) generate long sequence reads; average read lengths for the PacBio Single Molecule, Real-Time (SMRT) technology are 10–30 kilobases (9). These long reads have the potential to overcome many of the limitations of short read sequencing technologies including haplotyping and detection of structural variation. Indeed, SMS data has been successfully used for de novo assembly of human genomes (10; 11), identifying complex structural variation (12) and haplotype assembly of human genomes (13; 10). However, compared to short read sequencing technologies such as Illumina, the per-base accuracy of SMS reads is low with an error rate exceeding 10% (primarily due to insertion/deletion errors) (14). This high error rate makes the detection of small sequence variants such as SNVs, particularly heterozygous variants, difficult.

With the decreasing cost of SMS technologies and their increasing use for sequencing human genomes, accurate short variant calling methods for long read SMS data can be valuable in many ways. Current benchmarks for variant calling in human genomes, developed by the the Genome in a Bottle (GIAB) Consortium (15; 16), are based on short read sequence data and cover ~90.8% of the reference human genome sequence. These high-confidence variant calls are immensely valuable for developing new variant calling methods and sequencing technologies. However, these variant call sets are biased towards regions of the genome that are easy-to-call using short reads (17). Accurate SNV calling using long read SMS data can provide independent validation of short read SNV calls leading to reduction in false positives and increased understanding of systematic errors and artifacts. Furthermore, SNV calling using SMS reads can enable the generation of high-confidence variant calls in repetitive regions of the genome that include segmental duplications. The ability to call variants in repetitive regions that are inaccessible to short read sequencing technologies can also advance the use of SMS technologies for detection of disease causing mutations in duplicated genes via whole-genome or targeted sequencing (18).

Haplotype-resolved SNV detection from SMS reads can also enable the discovery of other types of human genetic variation, such as structural variants (SV) via separation of reads using haplotypes. Huddleston et al. (19) used an assembly-based approach, SMRT-SV, to identify thousands of SVs from whole-genome PacBio data of two haploid genomes, 89% of which were not reported by the 1000 Genomes Project (20). However, the sensitivity of SV detection using SMRT-SV was only 41% in diploid genomes. Chaisson et al performed dense whole-genome haplotyping of a human genome using multiple sequencing technologies, and were able to successfully call structural variants on each group of haplotype-separated SMS reads (21).

Variant calling tools such as GATK HaplotyperCaller (22) and Freebayes (23) developed for short read data analysis are not well-suited for SNV detection using PacBio data for two reasons: (i) short reads have low error rates (< 0.5%) and these methods do not model the high indel error rate of SMS reads which makes it difficult to distinguish true SNVs from errors and (ii) these methods analyze reads in short windows (typically a few hundred bases) and are not designed to leverage the haplotype information present in SMS reads. This haplotype information can be invaluable in distinguishing true variants from errors since observations of a true variant segregate with the reads originating from the haplotype it occurs on, whereas sequencing errors are unlikely to segregate. Recently, several methods for variant calling from long reads and deep-learning based variant calling methods have been developed (24; 25; 26). However, the accuracy of these methods for SNV calling on SMS data is currently much lower than that using Illumina whole-genome sequencing (WGS) (26; 25).

We describe a diploid SNV calling method, Longshot, that harnesses long SMS reads to jointly perform SNV detection and haplotyping. For this, it uses our read-based haplotype phasing method HapCUT2 (13). To overcome the high error rate of SMS reads, it utilizes a pair-Hidden Markov Model to average over the uncertainty in the local alignments and estimate accurate base quality values that can be used for calculating genotype likelihoods. We benchmarked Longshot using simulated data and whole-genome SMS data for multiple human individuals sequenced using the PacBio SMRT and Oxford Nanopore sequencing technologies (15; 16; 27).

## Results

### Overview of method

Alignments of SMS reads suffer from reference bias which can cause a SNV allele to be obscured by gaps (insertions and deletions) in the alignments (Supplementary Fig. S1A). Nevertheless, a true SNV is likely to have at least a few correctly aligned reads with the alternate allele. The first step in the Longshot algorithm identifies potential SNV sites using a standard pileup-based genotyping calculation (28) (Fig. 1A). A low variant quality threshold is used to select SNVs in order to minimize false negatives. Next, for each candidate SNV, we determine the most likely allele for each read covering the SNV and the corresponding estimate of the quality of the allele call (Fig. 1B). This allelotyping is done by local realignment of a segment of the read to short haplotype sequences (one for each of the two alleles at a biallelic SNV site). In low-complexity regions of the genome (e.g. homopolymers), there is significant ambiguity in the placement of gaps for SMS reads and many alignments are equally likely (29). Therefore, we use the forward algorithm on a sequence alignment pair-HMM (30) to perform the local realignment by averaging all possible local alignments of a read to a given haplotype.

**Figure 1:**
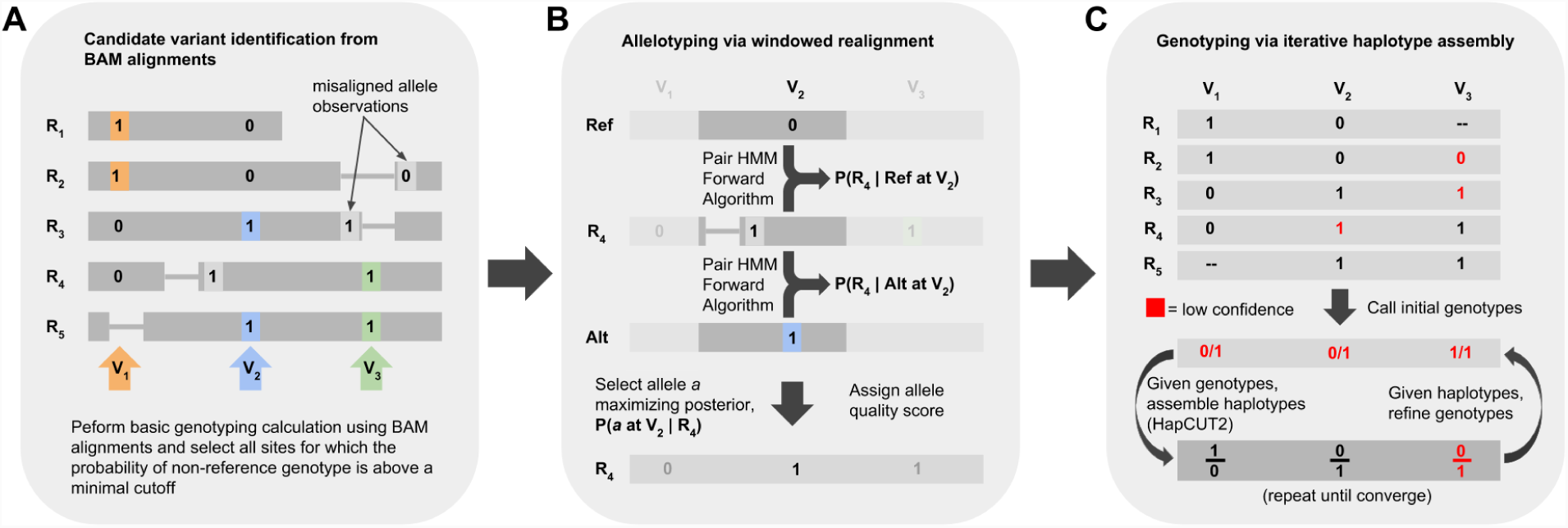
Overview of the Longshot algorithm. (A) Candidate variant identification. Candidate variants are identified using the pileup of the original alignments and a standard genotype likelihood calculation is used to determine if the site is a potential variant. A low threshold is used to maximize sensitivity. **(B) Allelotyping via local realignment.** To determine the allele for each read at each potential SNV site it overlaps, a window is formed around the variant and the probability of the observed read sequence given each allele is calculated using the forward algorithm on a Pair Hidden Markov Model. The most likely allele and quality score is chosen based on the relative likelihoods of the two alleles. **(C) Iterative phased genotyping.** Using the alleles and quality values for each read at variant sites, phased genotypes for all variants are determined jointly by performing haplotype assembly using HapCUT2 (on heterozygous variants) and local updates of the phased genotypes (individual sites) in an iterative manner.

After estimating the allele call and quality value for each read overlapping a SNV site, we estimate phased genotypes for all SNVs simultaneously using a haplotype-based likelihood model (see Methods). SMS reads typically cover multiple heterozygous sites and this haplotype information is useful since a SNV on a haplotype is expected to segregate with reads from the same haplotype (while random sequencing errors are not). In Longshot, heterozygous SNVs are assembled into haplotypes using HapCUT2 and a local update procedure is used to estimate the most likely phased genotype for each SNV given the current haplotypes for all other SNVs (Fig. 1C). This procedure is repeated for a few iterations until the likelihood stops improving. Finally, the variants are filtered for maximum read coverage, excessive variant density and minimum Genotype Quality (GQ) score, where the GQ score is estimated using the phased genotype likelihoods.

### Accurate SNV calling using simulated data

First, we used simulations to assess the accuracy of SNV calling using Longshot and also compared the precision and recall to short-read variant calling. We simulated a diploid genome by adding SNVs to the reference human genome, and simulated paired-end Illumina reads and PacBio SMS reads from this genome (maximum coverage of 60×). Subsequently, we aligned the reads to the reference genome using BWA-MEM (31) (Illumina) and BLASR (32)(PacBio) and called SNVs using FreeBayes and Longshot respectively. Across the entire genome, the precision was consistently high (≥0.9999) at all read coverages (20–60×) for both short read and SMS read-based SNV calling (Fig. 2). Short reads achieved greater recall than SMS reads at lower coverage (≤30×), while SMS reads had marginally greater recall at higher coverage (≥40×). SMS reads are expected to have better mappability in repetitive regions of the genome compared to Illumina reads, particularly in long segmental duplications with high sequence identity. Indeed, the recall for SMS reads in segmental duplications with high sequence similarity (≥95%, 127.5 Mb of DNA sequence) was significantly higher (0.86 at 40×coverage) compared to that using short reads (0.56 at 40×coverage) and increased with increasing coverage (Fig. 2).

**Figure 2:**
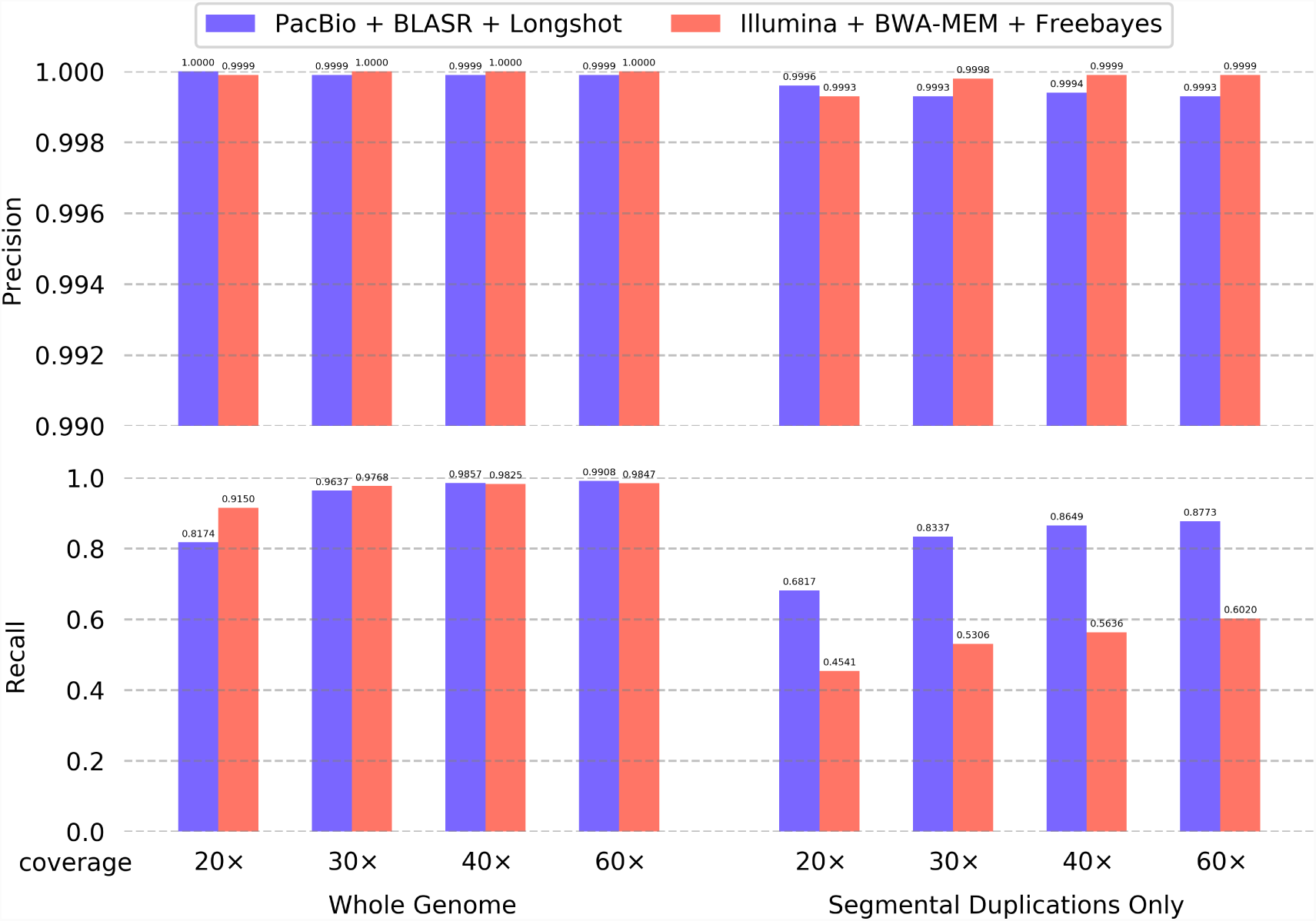
Accuracy of SNV calling using Longshot on simulated data and comparison with short-read based calling. Illumina and PacBio reads were generated at multiple coverages (20×, 30×, 40× and 60×) from a simulated diploid genome (chr1-22), mapped to the reference genome, and SNVs were called using Longshot (for PacBio) or Freebayes (for Illumina). Precision **(upper)** and Recall **(lower)** of the called variants were assessed across the whole genome **(left)** and within segmental duplications with > 95% similarity (**right)**.

We used the simulated datasets to estimate the theoretical fraction of the genome that is callable with SMS long reads compared to short reads at 60× coverage. We found that SMS reads were able to span 99.4% of the genome (non-N bases on chromosomes 1-22) with at least 30x coverage and at least 90% of well-mapped reads at each position (Supp. Fig S3). In comparison, Illumina reads covered 96.3% of the genome under these same criteria, a difference of 3.1%. We also compared the precision/recall of SNV calling using BLASR with several long-read mapping tools: NGMLR (33), BWA-MEM (31), and MINIMAP2 (34). All tools showed high precision and recall when considering SNVs across the whole genome, but BLASR had significantly higher recall (maximum of 0.88) than all other aligners (0.72 using Minimap2) in segmental duplications (Supplementary Fig. S2). Therefore, we utilized BLASR for the analysis of real datasets.

### Accurate haplotype-resolved SNV calling using whole-genome PacBio data

We used Longshot to call SNVs using whole-genome human PacBio data for four human genomes from the Genome in a Bottle (GIAB) consortium (15). Specifically, we used WGS data for the NA12878 individual (45×) and a mother-father-child trio of Ashkenazi ancestry (NA24385 at 64×, NA24149 at 29×, and NA24143 at 27×). For each dataset, a genotype quality threshold that was linearly proportional to the median read depth was used for filtering variants (see Methods). For comparison, we also called SNVs using Illumina short-read WGS data (~30×coverage) for each individual.

Longshot identified 3.51 to 3.65 million SNVs per genome and required an average of ~31 single core compute-hours for WGS with ~28× coverage (Supplementary Table S1). To assess the precision and recall of SNV calling, we utilized the GIAB high-confidence variant call set for each individual (15; 16). The comparison of SNV calls was limited to GIAB “confident regions” for each genome (16). The precision and recall for NA12878 were 0.9944 and 0.9583 respectively at 30× coverage and the recall improved to 0.9727 at 45× coverage (Figure 3). The precision and recall and the precision-recall curves (Supplementary Fig. S4) were highly consistent across the four genomes at 27-30× coverage (Figure 3), demonstrating the robustness of our method. To assess the improvement in precision/recall as a function of sequence coverage, we sub-sampled data for the AJ son individual (NA24385), who was sequenced to 64× coverage. The recall improved steadily from 0.9599 (28×) to 0.9792 (64×) while the precision only changed moderately with increasing coverage (0.9932 to 0.9937). The precision and recall for SNV calling using SMS reads was slightly lower than Illumina based variant calling (Figure 3). Nevertheless, the ability of Longshot to consistently achieve high recall (only 2-3% lower than Illumina WGS for the same depth of coverage) while achieving a low false discovery rate (average = 0.7%) was remarkable given the significantly high error rate of SMS reads (~10%) compared to Illumina reads.

**Figure 3:**
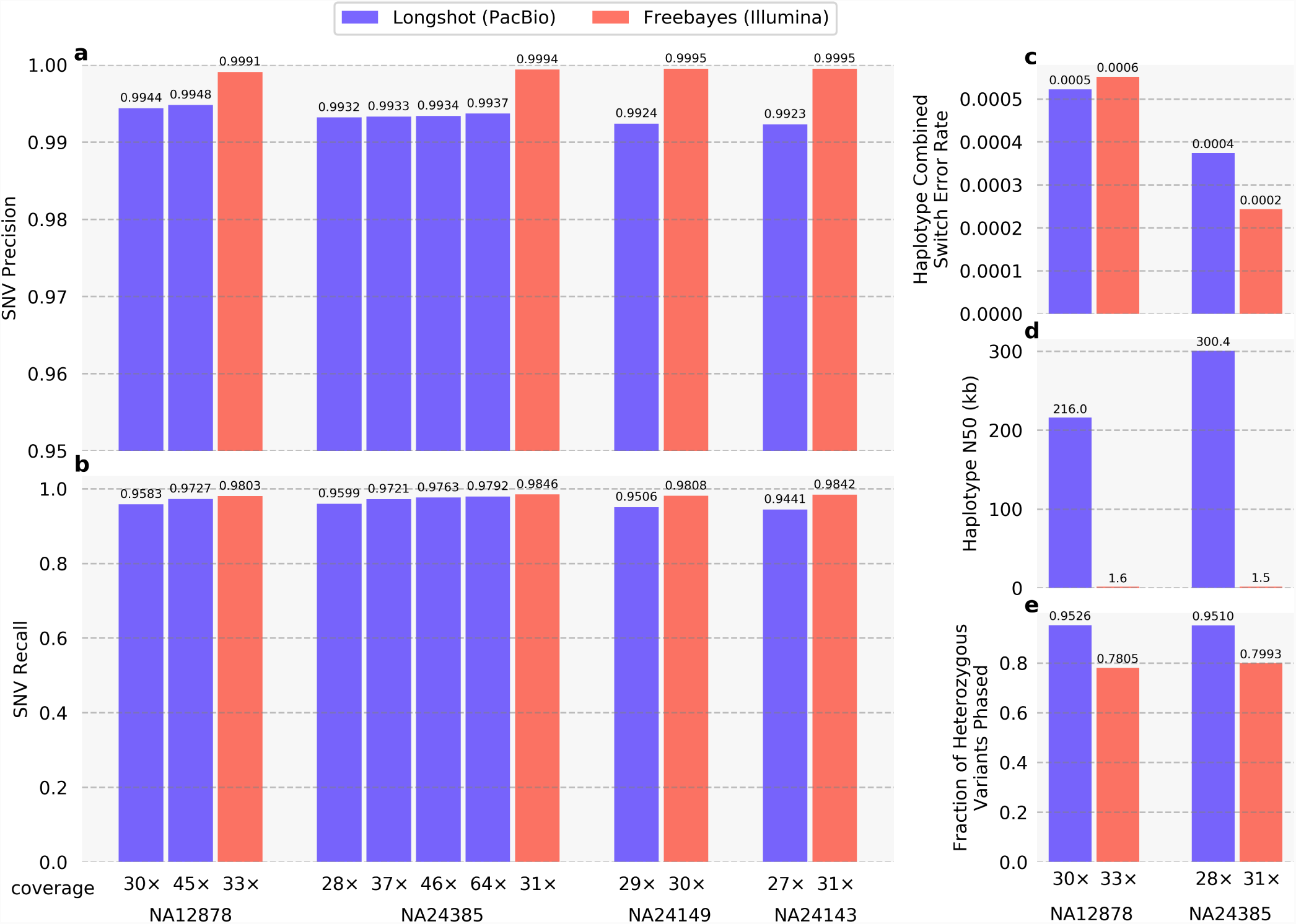
Accuracy and completeness of phased SNVs calls from Longshot using whole-genome SMS reads for four human individuals. Longshot was used to call SNVs using SMS data from the GIAB project for NA12878 (30× and 45× coverage), NA24385 (28×,37×,46×, and 64× coverage), NA24149 (29× coverage), and NA24143 (27× coverage). For each individual, variants were also called using Freebayes applied to ~30× coverage Illumina short reads for comparison. Precision **(a)** and recall **(b)** were calculated using GIAB variant calls as ground truth, limited to the GIAB confident regions. **(c)** The accuracy of Longshot SNV haplotypes was compared to those assembled for the short reads using HapCUT2 using the combined switch error rate (total rate of switch errors and mismatch errors). The completeness of the haplotypes is shown in terms of **(d)** N50 of haplotype blocks and **(e)** fraction of heterozygous variants phased. The median read lengths were 3,587 and 7,235 bp for NA12878 and NA24385, respectively.

In contrast with simulated data, the precision of of Longshot on real SMS reads was slightly lower than short read variant calling. To determine the source of false positive calls, we analyzed if such calls were enriched in specific sequence contexts or overlapped with indels. For the NA12878 dataset, (Table S2) we observed that the vast majority (71.8%) of false positive SNVs are located within 5 bp of a true indel, 190× the expected number. These false positive SNVs are called since the current Longshot implementation does not consider indels as potential variants. Filtering SNV calls located near known indels (using the “Mills + 1000 Genomes Gold Standard Indels” set from the GATK resource bundle (22)), reduced the number of false positives by 34-46% for the four GIAB genomes (table S3) while only slightly decreasing the recall. Analysis of false negative SNVs showed that 20.5% of false negative SNVs occurred inside homopolymer sequences of length 5 or greater, which is 3.5× the expected value. This follows naturally from the fact that these regions have low information content; insertion and deletion errors could plausibly lie anywhere along the length of a homopolymer. Therefore, allele calls inside homopolymers receive lower quality scores from the pair-HMM realignment, which reduces the power to call SNVs in such regions.

To compare Longshot’s accuracy on SMS data, we considered existing variant calling methods for short read data including GATK and FreeBayes. However, the GATK HC tool did not generate variant calls on the NA12878 PacBio dataset, consistent with previous evaluations of these methods on SMS data (25; 35). Recently, deep learning based methods for variant calling have been developed that can process short read data as well as SMS data (35; 25). On the PacBio dataset for NA12878 (from the GIAB project), identical to the one that we used for evaluating Longshot, the precision of LongShot was better than the reported precision for SNV calling of two recent deep learning based variant callers (25; 35) while the recall was virtually identical (Supp. Table S4). Ebler et al. have introduced a method that leverages the haplotype information of SMS reads in a HMM based model for accurate SNV detection and phasing (26). We were unable to directly evaluate this method since it currently requires a known set of variant sites as input (36). Nevertheless, comparison of the precision and recall of Longshot to the reported accuracy for this method (26), shows that Longshot identifies ~80% fewer false positives and ~33% fewer false negatives on the NA12878 PacBio dataset. Overall, Longshot is more accurate than existing methods for SNV calling (Supp. Table S4).

Unlike short-read variant calling, Longshot not only calls single nucleotide variants, but phases them into long haplotype blocks. We assessed the accuracy and completeness of haplotyping for two GIAB individuals, NA12878 and NA24385, by comparison to gold-standard haplotypes for these individuals inferred using pedigree data (see Methods). The Longshot haplotypes for NA12878 had an N50 length of 251.9 kb (with respect to the phased portion of the genome) and were very accurate, with a combined switch error rate of 0.05%. Similarly, the haplotypes for NA24385 had a N50 length 368.4 kb and a combined switch error rate equal to 0.04%. In comparison, haplotypes assembled using short reads had a N50 length less than 2 kb for both genomes (Figure 3). We also used HapCUT2 to assemble haplotypes for NA12878 and NA24385 using the SMS reads and SNVs identified using ~30× coverage Illumina sequencing (13). We found that the haplotype accuracy and completeness were very similar between Longshot and HapCUT2 (Supplementary Fig. S7). Separation of SMS reads using SNV haplotypes can enable discovery of non-SNV variants such as indels and structural variants using methods such as SMRT-SV that work well on haploid genomes. For the NA12878 dataset^1^, 50.9% of reads (weighted by length) could be assigned to a haplotype with high confidence. The ability to assign reads to haplotypes was correlated to read length: the haplotype-assigned reads had a median length of 4.3 kb while the unassigned reads had a median length of 2.6 kb.

### SNV calling using Oxford Nanopore reads

Recently, reads from Oxford Nanopore Technologies’ (ONT) MinION sequencer were used to assemble a human genome (27). Nanopore reads have a similar error profile to PacBio SMRT reads, however, the total per-base error rate of ONT reads is higher than for PacBio SMRT (37) and the errors are dependent on sequence context (38). We applied the Longshot algorithm to call SNVs using 37× coverage Oxford Nanopore reads (27) for a human individual (NA12878). We observed that the candidate set of SNVs considered by Longshot contained a significant fraction of false positives. This was due to the context-specific errors in Nanopore reads. To amelioriate this, we implemented a simple filter to remove potential SNVs for which the allele observations show a significant strand bias (Fisher’s exact test p-value < 0.01), prior to haplotype assembly. At a genotype quality threshold (GQ) of 60, Longshot achieved precision and recall values equal to 0.9628 and 0.8478 respectively (Supplementary Fig. S5). Similar to PacBio SMS data, the accuracy of Longshot on Oxford Nanopore data was significantly better than recent methods (Supplementary Table S4).

### Analysis of SNV calls in repetitive regions

As demonstrated with simulations, the recall of variant calling using SMS reads in segmental duplications with high sequence similarity (≥ 95%, Figure 2) is significantly higher (0.86) compared to short reads (0.56). These regions correspond to 102.8 Mb of the genome (excluding the sex chromosomes). However, 97.7% of these regions are excluded from the GIAB high-confidence variants, making it challenging to assess the accuracy of SNV calling using real SMS data. We compared SNV calls in segmental duplications for the NA12878 genome made using short read Illumina data (33×coverage) and SMS reads (30×coverage). In segmental duplications with ≥95% similarity, 180,584 SNVs were called using SMS reads, 54.8% more than those using Illumina reads (Table 1). The fewer calls using Illumina reads likely reflect the inability to map in segmental duplications. For example, Illumina reads cannot be mapped uniquely in a significant portion of the STRC gene, resulting in 52.3% fewer variants called compared to SMS reads (Fig. 4). The difference was more stark in segmental duplications with ≥ 99% similarity: 78,666 SNVs were called with SMS reads compared to only 18,684 with Illumina reads (4.2 fold difference). The Transition/Transversion (Ts/Tv) ratio for the SNVs called only using SMS reads in these regions was 1.99, slightly lower than the ratio for the SNV calls in GIAB confident regions (~2.1). This is consistent with the expectation that the Ts/Tv ratio is usually ~2.0-2.1 for SNVs across the whole genome (39). In contrast, the Ts/Tv ratio for Illumina-only calls in segmental duplications with ≥99% similarity was 1.54, much lower than the expected value (Table 1).

**Table 1:** Comparison of SNV calls using PacBio and Illumina WGS for the NA12878 genome. Variants were called using short reads (33× coverage) with Freebayes, and using SMS long reads (30× coverage) with Longshot. The number of variants called by each technology, the number of variants shared between the two technologies, and the corresponding transition/transversion (Ts/Tv) ratios are shown for the whole-genome and various subsets of the genome including GIAB high-confidence regions and segmental duplications with high sequence identity.

| | | Genome<br>(1-22) | Inside GIAB<br>Confident | Outside GIAB<br>Confident | Segmental Dup.<br>($\geq 95\%$ similar) | Segmental Dup.<br>($\geq 99\%$ similar) |
| --- | --- | --- | --- | --- | --- | --- |
| Region size |  | 2.8 Gb | 2.4 Gb | 330.7 Mb | 102.8 Mb | 47.5 Mb |
| PacBio | # SNVs | 3,514,112 | 3,000,524 | 513,588 | 180,584 | 78,666 |
|  | Ts/Tv | 2.08 | 2.14 | 1.75 | 1.95 | 1.99 |
| Illumina | # SNVs | 3,563,787 | 3,065,573 | 498,214 | 116,649 | 18,684 |
|  | Ts/Tv | 2.03 | 2.1 | 1.66 | 1.84 | 1.79 |
| Unique to PacBio | # SNVs | 252,777 | 63,700 | 189,077 | 103,383 | 69,507 |
|  | Ts/Tv | 1.62 | 1.82 | 1.57 | 1.9 | 1.99 |
| Unique to Illumina | # SNVs | 302,493 | 128,751 | 173,742 | 39,464 | 9,525 |
|  | Ts/Tv | 1.29 | 1.25 | 1.33 | 1.53 | 1.56 |
| Shared<br>Illumina & PacBio | # SNVs | 3,261,323 | 2,936,824 | 324,499 | 77,186 | 9,159 |
|  | Ts/Tv | 2.12 | 2.15 | 1.86 | 2.0 | 2.03 |

**Figure 4:**
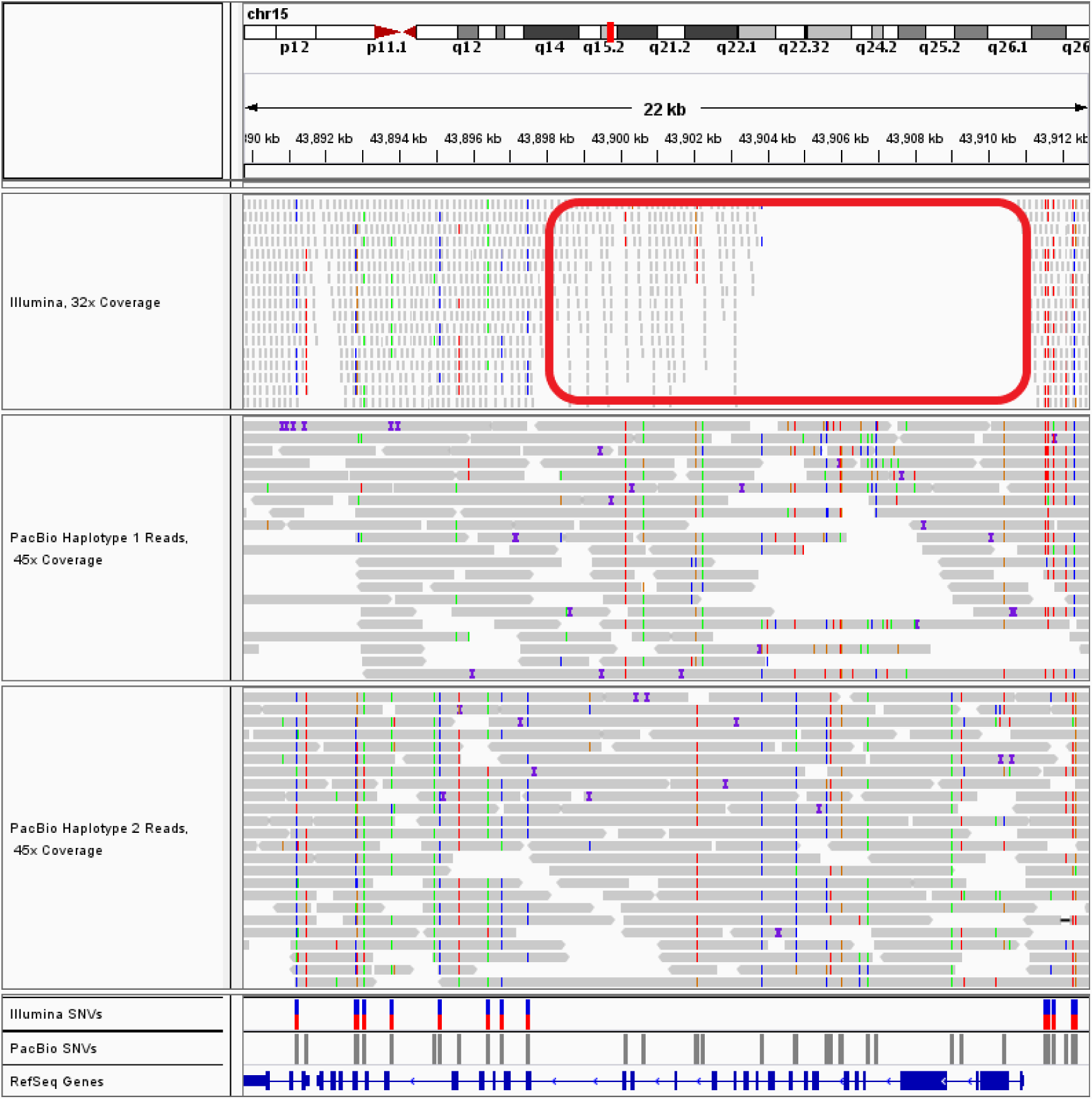
Accurate SNV calling using SMS reads and Longshot in the duplicated gene STRC. An Integrated Genomics Viewer (IGV) view of mapped reads shows that a long segment of the gene (highlighted in red) has low coverage using uniquely mapped Illumina reads due to the presence of a long segmental duplication with high sequence similarity (> 99.8%) that spans the entire gene. PacBio SMS reads (separated by haplotype using Longshot phased SNVs) have consistent coverage of mapped reads across the entire gene, allowing Longshot to call 40 SNVs of which 20 variants are shared with short reads, and 22 variants are unique to Longshot.

Next, we assessed the Mendelian consistency of SNV calls for the mother-father-child trio of Ashkenazi ancestry from the GIAB project. To minimize discordance due to false negative calls, only sites with at least 20× read coverage in every individual were considered. SMS calls in the high confidence GIAB regions had higher concordancy (98.87%) compared to calls outside GIAB confident regions (96.13%). Within segmental duplications (≥ 95% similarity), 5.00% of the SNVs in the child were discordant with Mendelian inheritance. Many of the discordant SNVs were clustered in contiguous blocks, indicating that they are the result of mismapped reads or structural variation in one or more individuals.

Finally, we compared Longshot SNV calls for NA12878 to the Platinum Genomes small variant call set for this genome that have been generated using Illumina WGS and validated using haplotype inheritance on a 17-member pedigree (40). In GIAB high confidence regions, 97.7% of the PG SNVs were also called by Longshot. The PG calls cover a significant fraction of the genome (330.7 Mb) that is excluded from the GIAB high-confidence calls. In these regions, only 76.8% of the PG SNVs were shared with Longshot and 75,061 SNVs were unique to the PG calls (Fig. S8). The low concordance in regions outside the GIAB high-confidence regions highlights the challenge of accurate variant calling in these regions. Longshot’s ability to call SNVs accurately using SMS reads provides an orthogonal validation for SNVs called using short reads. In-depth analysis of variant calls made using short-read and SMS data in these regions can enable the expansion of confidently called regions for reference human genomes.

## Discussion

Our results demonstrate that highly accurate detection of SNVs is feasible even from long-read sequence data with high error rates. Combined with recent work demonstrating the ability to detect and genotype structural variants from SMRT-seq data, our results indicate that long-read sequencing can be used to accurately detect all forms of genetic variation in human genomes. Recently, Li et al wrote that “although PacBio assembly is accurate at the base-pair level for haploid genomes, it is currently not accurate enough to confidently call heterozygotes in diploid mammalian genomes” (17). We have demonstrated that heterozygous SNVs can be called accurately in diploid genomes, by combining sensitive allelotyping of reads at SNV sites with haplotype-informed genotyping. Our method achieves a precision greater than 99.1% across multiple whole-genome PacBio datasets. Furthermore, we have demonstrated that even higher precision (99.7%) can be achieved by filtering out known common indels. Moreover, the GIAB and Platinum Genomes variant sets used to assess accuracy were both generated using short read datasets, are therefore biased in favor of short-read technologies (17) and exclude regions where long reads are likely to have better precision and recall. Therefore, the true accuracy for Longshot is likely even better.

We have also demonstrated that SMS reads can be used to call SNVs in segmental duplications and other regions of the genome with low short-read mappability. However, correctly mapping PacBio reads in highly similar segmental duplications remains a challenge. As Supplementary Fig. S2 shows, there is wide variance in the ability of SMS read mappers to map reads in segmental duplications. This is likely due to the mappers having different strategies for dealing with highly similar potential mappings that are only different by some number of paralog-specific-variants (PSVs). Despite BLASR performing relatively well using simulated reads, many of the discordant SNVs observed between the AJ trio in segmental duplications appeared to be caused by the presence of multiple mismapped reads. SMS read mapping methods with specific optimizations for segmental duplications could alleviate this (34).

Longshot offers the ability to assemble haplotypes without prior knowledge of SNVs and leverage the haplotypes to separate SMS reads by haplotype. This opens up a wide range of possibilities for SMS read analysis, given that many SMS analysis tools work much better on haploid samples. For example, the haplotype-separated reads could be used to call structural variants with greater sensitivity using a tool such as SMRT-SV (11). A similar approach was recently used to profile structural variation genome-wide after extensive haplotype assembly with multiple sequencing technologies and computational separation of the SMS reads by haplotype (21). Similarly, new variants could be uncovered by generating a genomic consensus of the haplotype-separated read sets using tools such as Arrow (41). Currently, Longshot uses the read pileups to identify candidate SNVs and the vast majority (~72%) of false positive SNVs identified with Longshot correspond to misclassified indel variants. Using a genomic consensus of haplotype separated reads should improve the accuracy of variant calling using LongShot.

In this paper, we focused on the detection and phasing of single nucleotide variants alone since accurate calling of short indels using SMS reads is challenging due to the high insertion/deletion error rate. A recently developed deep-learning based variant caller (25) had low precision (0.589) and recall (0.12) for short indel calling on PacBio WGS data. PacBio circular consensus sequencing (CCS) produces reads with greater accuracy by sequencing multiple times around the same DNA template. Recent improvements to the PacBio technology have enabled the generation of highly accurate long reads (10-15 kilobases read lengths and error rates < 1%) using CCS (42). We expect that using Longshot with these accurate reads will improve the accuracy of SNV calling and also enable accurate short indel calling.

Finally, our method was able to call SNVs using Oxford Nanopore sequencing reads with very good accuracy without any modification to the likelihood model. Although the precision and recall was significantly lower than PacBio reads at similar coverage, this is expected due to the higher error rate of Nanopore reads. It may be feasible to improve the precision and recall using Nanopore WGS data but this will require the development of context-specific error models for local realignment and use of consensus based methods for identifying variants. Continued improvements in the raw accuracy of Nanopore reads and availability of additional high-coverage whole-genome Nanopore datasets will also facilitate accurate small variant calling.

## Methods

### Longshot Algorithm

#### (i) Identification of candidate SNVs

The first step in the Longshot algorithm is to identify positions in the genome that may contain a SNV. Potential SNVs are identified from the pileup of aligned reads by performing a genotype likelihood calculation similar to Samtools or other NGS variant calling methods (3) (see Supplementary methods). The prior probabilities for genotypes are defined using a slight modification of the approach of Li et al (43) (see Supplemental Methods). SNV sites for which the posterior probability of a non-reference genotype is greater than 0.01 are considered as candidate SNVs for the next step of the algorithm. The sites are also filtered for mininum read coverage (6 by default), minimum alternate allele count and fraction (3 and 0.125 by default).

#### (ii) Local realignment using pair-HMMs

For an SMS read that overlaps a candidate bi-allelic SNV site with two alleles “ref” and “alt”, we want to determine which allele is the most likely observation (allele call) and also assign a probability of error to this observation (quality value). To accomplish this, we perform local realignment of a short sequence from the read to the reference and to the alternate sequence (with the SNV allele added, see Fig 1B). This local realignment is performed using pair-Hidden Markov Model (pair-HMM) (30). The parameters for the HMM are estimated directly from the aligned reads prior to realignment (see Supplementary Methods).

It is sufficient to perform the local realignment within a short window covering the SNV site. This window is defined using the nearest non-repetitive “anchor sequences” of length *k* (default *k* = 6), to the left and right of the SNV where the read sequence matches the reference sequence perfectly (see Supplemental Methods). Once the window *W* is identified, we use the forward algorithm to calculate *p*_ref_ = *P* (read(W) | ref(W)) and *p*_alt_ = *P* (read(W) | alt(W)) where read(*W*) is the sequence of the read in the window *W* defined by the two anchors. We select the allele *a*_*max*_ ∈ {ref, alt} for which the probability is higher, as the observed allele and use phred 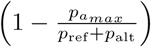 as the allele quality score.

When multiple candidate SNVs are located in close proximity, we define the window to include all such SNVs and use a generalization of the calculation described above to determine alleles and estimate base quality values (see Supplemental Methods). For computational efficiency, a “banded” version of the forward algorithm is used. This reduces the complexity to *O*(*mb*) where *b* is the width of the band and *m* is the length of the window (50-200 bp). Allele observations with phred-scaled quality score below a threshold (7.0 by default) are discarded. This reduces the effective read depth for SMS reads (Supp. Fig. S6).

In order to remove false variants resulting from strand-specific sequencing errors, we filter potential SNVs whose allele observations are over-represented in reads from one strand. For each potential SNV, we build a contingency table of the counts of the reference and alternate allele on reads from the forward strand and reverse strand respectively. Variants for which the Fisher’s exact test p-value (2-tailed) is less than 0.01 are not considered for haplotype-informed genotyping.

#### (iii) Haplotype-informed genotyping

Longshot achieves accurate variant calling using SMS reads by performing phased genotyping for all candidate SNVs jointly. Given a set of candidate SNVs *V* and the allele calls (and quality values) for each read *r* ∈ *R*, we aim to maximize the likelihood *p*(*R*|*H*) where *H* is a pair of haplotypes (*H*_1_, *H*_2_) over the variant set *V*. Longshot optimizes the likelihood function using an iterative approach that uses (i) the HapCUT2 algorithm (13) to estimate the most likely pair of haplotypes for variants with heterozygous genotypes and (ii) local updates to estimate the most likely phased genotype for each variant given the current haplotype pair (Fig. 1C).

Assuming independence between reads, the likelihood function *p*(*R*|*H*) can we written as (13):

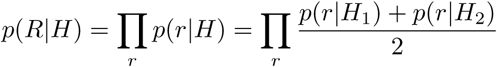

*p*(*r*|*H*_1_) for any read *r* can be calculated using the pair-HMM probabilities for each (read,variant) pair. Let *G* be the set of possible phased genotypes for a biallelic variant: {0|0, 0|1, 1|0, 1|1} (homozygous reference, the two heterozygous states, and homozygous alternate). Let *H* refer to the current estimate of the most likely haplotype pair, and *H*^*i,g*^ refer to the haplotype pair *H* with the *i*th SNV altered to have the phased genotype *g*. Given *H*, we can calculate the posterior probability for the phased genotype *g* as follows:

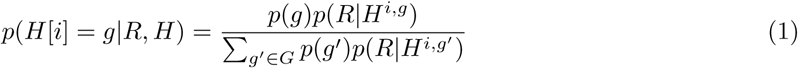

The optimization starts with the initial set of variants identified from the pileup-based likelihood calculation and the unphased genotypes for each variant estimated using the local realignment. The iterative phase of the algorithm consists of the following steps:

1. *L* = *p*(*R*|*H*)
2. *H* = HapCUT2(*R, V*)
3. Repeat:
  a. For each variant *v* ∈ *V*: update *H*[*v*] using equation 1
  b. If no genotype was updated in (a), BREAK
4. *L′* = *p*(*R*|*H*)

In Step 2, HapCUT2 is used to phase only heterozygous variants. Steps 1-4 are repeated until the relative log-likelihood of the data between consecutive iterations is smaller than Δ (default = 1 × 10^-5^).

### Variant filtering

The raw variant calls were subjected to three types of filters to reduce false positives. SNVs were first filtered according by genotype quality (GQ) estimated by the variant caller. The GQ cutoff was fixed at 50 for short reads. For Longshot, we used a variable GQ cutoff (matched to the median read coverage) for filtering variants. This was done to reduce the number of false SNVs due to true indel variants that have high GQ. For simulations, which do not have indel variants, we used a fixed genotype quality of cutoff of 50.

To filter false SNVs due to copy number amplifications, a maximum read depth filter similar to what has been used previously for short-read based variant calling was used (44). Variants with read depth greater than 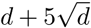, where *d* is the median read depth across the entire dataset were filtered out. We also observed that for SMS reads, many false positive SNVs occur nearby each other in dense clusters. These dense clusters may result from systematically mismapped reads due to missing sequence in the reference genome, or are indicative of structural variations such as CNVs. We used a simple density filter (> 10 SNVs in a window of 500 base pairs) to filter out such false variants for variants called with Longshot. For the AJ trio, variants in the delly exclusion regions (notably centromeres) were also filtered out for the analysis of variant mendelian consistency (45).

### Simulations

We simulated a diploid genome (chromosomes 1-22) with heterozygous SNVs (rate = 0.001) and homozygous SNVs (rate = 0.0005) as described previously (43) (see Supp. Methods for details). Paired-end 100bp reads were generated from the simulated genome with a substitution error rate of 0.001 (46). The short reads were aligned to the human reference (hs37d5) using BWA-MEM, and variants were called using Freebayes (23). Similarly, we used SimLoRD (47) to generate PacBio SMS reads (median length = 7.5 kb) from the simulated genome using the default error rates of 0.11 for insertion, 0.04 for deletion, and 0.01 for substitution (47). The `-mp 1` option was used to force each read to only have a single sequencing pass, so that the error profile of the reads resembles PacBio “continuous long reads” (lower accuracy) as opposed to “circular consensus reads” (greater accuracy). We aligned the SMS reads to the human reference (hs37d5) using the long-read alignment tools BLASR (`v5.3.2`, options `--nproc 16 --bestn 1 --bam`), MiniMap2 (`v2.11-r797`, options `-t 16 -ax map-pb`), BWA-MEM (`v0.7.12-r1044`, options `-x pacbio -t 16 -T 0`) and NGMLR (`v0.2.7`, options `-t 16 -x pacbio`).

### Whole-Genome PacBio, Oxford Nanopore, and Illumina data

45× coverage Pacific Biosciences Single Molecule Real Time (SMRT) reads for NA12878, aligned to the hs37d5 reference genome using BLASR, were obtained from the Genome in a Bottle consortium (48). PacBio read data for the AJ trio was also obtained from the GIAB ftp site and aligned to the hg38 reference genome using BLASR (32), using the same parameters used for aligning the simulated reads. 37× coverage Oxford Nanopore reads for NA12878, aligned to hg38 using minimap2, were obtained from the Nanopore WGS Consortium (27). Illumina WGS data for NA12878 and the AJ Trio (NA24385, NA24143, NA24149), sequenced on the HiSeq 2500 (30× and 60× coverage respectively, 148 bp paired-end reads), was obtained from the GIAB. The 60× coverage datasets were downsampled to half coverage. The reads were downloaded in BAM format aligned to as hs37d5 using bwa-mem (NA12878) and hg38 using NovoAlign (AJ trio) (49). Variant calling on Illumina WGS data was performed using Freebayes (23) (`v1.0.2-33-gdbb6160)` with `--standard-filters` and `--genotype-qualities` turned on). BED files for segmental duplications and repeat elements in the human genome were obtained from the UCSC table browser (50).

### Assessment of variant calling and phasing accuracy

High confidence variant call sets generated by the GIAB project were used for assessing accuracy of SNV calling (48; 16). For NA12878, SNVs were compared against the GrCh37 (for Illumina and PacBio) or GrCh38 (for Oxford Nanopore) version of the GIAB high-confidence call set (release v3.3.2). For the AJ trio, SNVs were compared against the GrCh38 version of the GIAB high-confidence call set (release v3.3.2). For each individual, the comparison of SNV calls was limited to “confident regions” (provided in a bed file). Precision and Recall were calculated using RTGtools (v3.9) vcfeval.

For NA12878, we compared the accuracy of the Longshot haplotypes using the Platinum Genomes haplotypes for the same individual as ground truth. For NA24385, we generated high-quality haplotypes from a consensus of the GIAB trio-based phased genotypes and 10X Genomics phased variant calls and used the resulting haplotypes for assessment of haplotyping accuracy. The haplotypes were compared at all unfiltered SNVs that were called heterozygous in both the assembled haplotypes and the ground truth. The errors were tabulated in terms of the total combined rate of “switch” and “mismatch” errors, also known as “long switch” and “short switch” errors respectively (13; 51). The N50 metric was used to measure the completeness of haplotype blocks, defined as the length ‘N’ in base pairs such that half of the phased portion of the genome is in haplotype blocks of length N or greater (6).

For the AJ trio, Mendelian consistency of the SNV calls was assessed using RTGtools (52). For this, SAMtools (28) and BEDtools (53) were used to obtain a set of regions that have high coverage (> 20×) of well-mapped SMS reads (*MAPQ >* 30 and filter `-F 3844` applied) in all three individuals. These regions were further intersected with a bed file for the region being investigated (either GIAB confident regions, outside GIAB confident regions, or 95% similar segmental duplications). The individual VCFs for the trio were merged into a single VCF and filtered so that all records have a genotype quality greater than 50.

### Implementation and software availability

Longshot is implemented in the Rust programming language, uses the rust-bio and rust-htslib libraries (54) and the HapCUT2 C code (13), and is available for download at https://github.com/pjedge/longshot. A Snakemake workflow (55) for automatically generating all of the results of the paper is available at https://github.com/pjedge/longshot_study.

## Data Availability

The PacBio and Illumina sequence datasets and variant calls used in this paper are publicly available from the GIAB ftp site: ftp://ftp-trace.ncbi.nlm.nih.gov/giab/ftp/. The sub-folders for the Illumina reads, VCF calls and Illumina reads for each individual are as follows:

- **NA12878**: data/NA12878/NA12878 PacBio MtSinai, release/NA12878 HG001/NISTv3.3.2/GRCh37/, data/NA12878/NIST NA12878 HG001 HiSeq 300x/
- **NA24385**: data/AshkenazimTrio/HG002 NA24385 son/PacBio MtSinai NIST, release/AshkenazimTrio/HG002 NA24385 son/NISTv3.3.2/GRCh38, data/AshkenazimTrio/HG002 NA24385 son/NIST HiSeq HG002 Homogeneity-10953946
- **NA24149**: data/AshkenazimTrio/HG003 NA24149 father/PacBio MtSinai NIST/, release/AshkenazimTrio/HG003 NA24149 father/NISTv3.3.2/GRCh38, data/AshkenazimTrio/HG003 NA24149 father/NIST HiSeq HG003 Homogeneity-12389378
- **NA24143**: /data/AshkenazimTrio/HG003 NA24143 mother/PacBio MtSinai NIST, release/AshkenazimTrio/HG004 NA24143 mother/NISTv3.3.2/GRCh38, data/AshkenazimTrio/HG004 NA24143 mother/NIST HiSeq HG004 Homogeneity-14572558

The Oxford Nanopore sequence dataset is publicly available from the Nanopore WGS Consortium: https://s3.amazonaws.com/nanopore-human-wgs/rel5-guppy-0.3.0-chunk10k.sorted.bam

## Acknowledgments

The research was supported by the National Human Genome Research Institute of the National Institute of Health (award number R01HG010149).

## Author Contributions

P.E. designed and implemented the algorithm, performed the analyses and wrote the manuscript. V.B. conceived the project, designed the algorithm, performed analyses and wrote the manuscript.

## Competing Financial Interests

The authors declare no competing financial interests.

## Footnotes

1 chromosome 1 only

## References

[1] S. Goodwin, J. D. McPherson, and W. R. McCombie, “Coming of age: ten years of next-generation sequencing technologies,” Nat. Rev. Genet., vol. 17, no. 6, pp. 333–351, 05 2016.

[2] D. Sims, I. Sudbery, N. E. Ilott, A. Heger, and C. P. Ponting, “Sequencing depth and coverage: key considerations in genomic analyses,” Nature Reviews Genetics, vol. 15, no. 2, p. 121, 2014.

[3] R. Nielsen, J. S. Paul, A. Albrechtsen, and Y. S. Song, “Genotype and SNP calling from next-generation sequencing data,” Nat. Rev. Genet., vol. 12, no. 6, pp. 443–451, Jun 2011.

[4] V. Bansal and V. Bafna, “HapCUT: an efficient and accurate algorithm for the haplotype assembly problem,” Bioinformatics, vol. 24, no. 16, pp. i153–159, Aug 2008.

[5] S. Levy, G. Sutton, P. Ng, L. Feuk, A. Halpern, B. Walenz, N. Axelrod, J. Huang, E. Kirkness, G. Denisov, Y. Lin et al., “The diploid genome sequence of an individual human,” PLoS Biol., vol. 5, p. e254, Sep 2007.

[6] J. Duitama, G. K. McEwen, T. Huebsch, S. Palczewski, S. Schulz, K. Verstrepen, E. K. Suk, and M. R. Hoehe, “Fosmid-based whole genome haplotyping of a HapMap trio child: evaluation of Single Individual Haplotyping techniques,” Nucleic Acids Res., vol. 40, no. 5, pp. 2041–2053, Mar 2012.

[7] K. Bryc, N. Patterson, and D. Reich, “A novel approach to estimating heterozygosity from low-coverage genome sequence,” Genetics, vol. 195, no. 2, pp. 553–561, Oct 2013.

[8] D. Mandelker, R. J. Schmidt, A. Ankala, K. McDonald Gibson, M. Bowser, H. Sharma, E. Duffy, M. Hegde, A. Santani, M. Lebo, and B. Funke, “Navigating highly homologous genes in a molecular diagnostic setting: a resource for clinical next-generation sequencing,” Genet. Med., vol. 18, no. 12, pp. 1282–1289, 12 2016.

[9] “Smrt sequencing: Read lengths,” https://www.pacb.com/smrt-science/smrt-sequencing/read-lengths, accessed: 2018-10-04.

[10] M. Pendleton, R. Sebra, A. W. Pang, A. Ummat, O. Franzen, T. Rausch, A. M. Stutz, W. Stedman, T. Anantharaman, A. Hastie et al., “Assembly and diploid architecture of an individual human genome via single-molecule technologies,” Nat. Methods, vol. 12, no. 8, pp. 780–786, Aug 2015.

[11] M. J. Chaisson, J. Huddleston, M. Y. Dennis, P. H. Sudmant, M. Malig, F. Hormozdiari, F. Antonacci, U. Surti, R. Sandstrom, M. Boitano et al., “Resolving the complexity of the human genome using single-molecule sequencing,” Nature, vol. 517, no. 7536, p. 608, 2015.

[12] M. J. Chaisson, S. Mukherjee, S. Kannan, and E. E. Eichler, “Resolving multicopy duplications de novo using polyploid phasing,” in International Conference on Research in Computational Molecular Biology. Springer, 2017, pp. 117–133.

[13] P. Edge, V. Bafna, and V. Bansal, “Hapcut2: robust and accurate haplotype assembly for diverse sequencing technologies,” Genome research, vol. 27, no. 5, pp. 801–812, 2017.

[14] S. Ardui, A. Ameur, J. R. Vermeesch, and M. S. Hestand, “Single molecule real-time (SMRT) sequencing comes of age: applications and utilities for medical diagnostics,” Nucleic Acids Res., vol. 46, no. 5, pp. 2159–2168, Mar 2018.

[15] J. M. Zook, B. Chapman, J. Wang, D. Mittelman, O. Hofmann, W. Hide, and M. Salit, “Integrating human sequence data sets provides a resource of benchmark snp and indel genotype calls,” Nature biotechnology, vol. 32, no. 3, p. 246, 2014.

[16] J. Zook, J. McDaniel, H. Parikh, H. Heaton, S. A. Irvine, L. Trigg, R. Truty, C. Y. McLean, F. M. De La Vega, M. Salit et al., “Reproducible integration of multiple sequencing datasets to form high-confidence snp, indel, and reference calls for five human genome reference materials,” bioRxiv, p. 281006, 2018.

[17] H. Li, J. M. Bloom, Y. Farjoun, M. Fleharty, L. Gauthier, B. Neale, and D. MacArthur, “A synthetic-diploid benchmark for accurate variant-calling evaluation,” Nature methods, vol. 15, no. 8, p. 595, 2018.

[18] D. M. Borras, R. H. A. M. Vossen, M. Liem, H. P. J. Buermans, H. Dauwerse, D. van Heusden, R. T. Gansevoort, J. T. den Dunnen, B. Janssen, D. J. M. Peters, M. Losekoot, and S. Y. Anvar, “Detecting PKD1 variants in polycystic kidney disease patients by single-molecule long-read sequencing,” Hum. Mutat., vol. 38, no. 7, pp. 870–879, 07 2017.

[19] J. Huddleston, M. J. Chaisson, K. M. Steinberg, W. Warren, K. Hoekzema, D. Gordon, T. A. Graves-Lindsay, K. M. Munson, Z. N. Kronenberg, L. Vives et al., “Discovery and genotyping of structural variation from long-read haploid genome sequence data,” Genome research, 2016.

[20] P. H. Sudmant, T. Rausch, E. J. Gardner, R. E. Handsaker, A. Abyzov, J. Huddleston, Y. Zhang, K. Ye, G. Jun, M. H.-Y. Fritz et al., “An integrated map of structural variation in 2,504 human genomes,” Nature, vol. 526, no. 7571, p. 75, 2015.

[21] M. J. Chaisson, A. D. Sanders, X. Zhao, A. Malhotra, D. Porubsky, T. Rausch, E. J. Gardner, O. Rodriguez, L. Guo, R. L. Collins et al., “Multi-platform discovery of haplotype-resolved structural variation in human genomes,” bioRxiv, p. 193144, 2018.

[22] A. McKenna, M. Hanna, E. Banks, A. Sivachenko, K. Cibulskis, A. Kernytsky, K. Garimella, D. Altshuler, S. Gabriel, M. Daly et al., “The genome analysis toolkit: a mapreduce framework for analyzing next-generation dna sequencing data,” Genome research, 2010.

[23] E. Garrison and G. Marth, “Haplotype-based variant detection from short-read sequencing,” arXiv preprint arXiv:1207.3907, 2012.

[24] F. Guo, D. Wang, and L. Wang, “Progressive approach for snp calling and haplotype assembly using single molecular sequencing data,” Bioinformatics, vol. 34, no. 12, pp. 2012–2018, 2018.

[25] R. Poplin, P. C. Chang, D. Alexander, S. Schwartz, T. Colthurst, A. Ku, D. Newburger, J. Dijamco, N. Nguyen, P. T. Afshar, S. S. Gross, L. Dorfman, C. Y. McLean, and M. A. DePristo, “A universal SNP and small-indel variant caller using deep neural networks,” Nat. Biotechnol., vol. 36, no. 10, pp. 983–987, Nov 2018.

[26] J. Ebler, M. Haukness, T. Pesout, T. Marschall, and B. Paten, “Haplotype-aware genotyping from noisy long reads,” bioRxiv, p. 293944, 2018.

[27] M. Jain, S. Koren, K. H. Miga, J. Quick, A. C. Rand, T. A. Sasani, J. R. Tyson, A. D. Beggs, A. T. Dilthey, I. T. Fiddes et al., “Nanopore sequencing and assembly of a human genome with ultra-long reads,” Nature biotechnology, vol. 36, no. 4, p. 338, 2018.

[28] H. Li, B. Handsaker, A. Wysoker, T. Fennell, J. Ruan, N. Homer, G. Marth, G. Abecasis, and R. Durbin, “The sequence alignment/map format and samtools,” Bioinformatics, vol. 25, no. 16, pp. 2078–2079, 2009.

[29] M. Jain, I. T. Fiddes, K. H. Miga, H. E. Olsen, B. Paten, and M. Akeson, “Improved data analysis for the MinION nanopore sequencer,” Nat. Methods, vol. 12, no. 4, pp. 351–356, Apr 2015.

[30] R. Durbin, S. R. Eddy, A. Krogh, and G. Mitchison, Biological sequence analysis: probabilistic models of proteins and nucleic acids. Cambridge university press, 1998.

[31] H. Li, “Aligning sequence reads, clone sequences and assembly contigs with bwa-mem,” arXiv preprint arXiv:1303.3997, 2013.

[32] M. J. Chaisson and G. Tesler, “Mapping single molecule sequencing reads using basic local alignment with successive refinement (blasr): application and theory,” BMC bioinformatics, vol. 13, no. 1, p. 238, 2012.

[33] F. J. Sedlazeck, P. Rescheneder, M. Smolka, H. Fang, M. Nattestad, A. von Haeseler, and M. C. Schatz, “Accurate detection of complex structural variations using single molecule sequencing,” Preprint at https://www.biorxiv.org/content/arly/2017/07/28/169557, 2017.

[34] H. Li, “Minimap2: fast pairwise alignment for long nucleotide sequences,” ArXiv e-prints [Internet], 2017.

[35] R. Luo, F. J. Sedlazeck, T.-W. Lam, and M. Schatz, “Clairvoyante: a multi-task convolutional deep neural network for variant calling in single molecule sequencing,” bioRxiv, p. 310458, 2018.

[36] “Whatshap,” https://bitbucket.org/whatshap/whatshap, accessed: 2018-10-22.

[37] E. Karlsson, A. Lärkeryd, A. Sjödin, M. Forsman, and P. Stenberg, “Scaffolding of a bacterial genome using minion nanopore sequencing,” Scientific reports, vol. 5, p. 11996, 2015.

[38] N. J. Loman, J. Quick, and J. T. Simpson, “A complete bacterial genome assembled de novo using only nanopore sequencing data,” Nat. Methods, vol. 12, no. 8, pp. 733–735, Aug 2015.

[39] M. A. DePristo, E. Banks, R. Poplin, K. V. Garimella, J. R. Maguire, C. Hartl, A. A. Philippakis, G. Del Angel, M. A. Rivas, M. Hanna et al., “A framework for variation discovery and genotyping using next-generation dna sequencing data,” Nature genetics, vol. 43, no. 5, p. 491, 2011.

[40] M. A. Eberle, E. Fritzilas, P. Krusche, M. Källberg, B. L. Moore, M. A. Bekritsky, Z. Iqbal, H.-Y. Chuang, S. J. Humphray, A. L. Halpern et al., “A reference data set of 5.4 million phased human variants validated by genetic inheritance from sequencing a three-generation 17-member pedigree,” Genome research, vol. 27, no. 1, pp. 157–164, 2017.

[41] “Genomicconsensus,” https://github.com/PacificBiosciences/GenomicConsensus, accessed: 2018-10-22.

[42] “High-fidelity 15kb long read dataset of hg002, ashkenazim son,” ftp://ftp-trace.ncbi.nlm.nih.gov/giab/ftp/data/AshkenazimTrio/HG002NA24385son/PacBioCCS15kb/, accessed: 2018-10-24.

[43] R. Li, Y. Li, X. Fang, H. Yang, J. Wang, K. Kristiansen, and J. Wang, “Snp detection for massively parallel whole-genome resequencing,” Genome research, vol. 19, no. 6, pp. 1124–1132, 2009.

[44] H. Li, “Toward better understanding of artifacts in variant calling from high-coverage samples,”Bioinformatics, vol. 30, no. 20, pp. 2843–2851, Oct 2014.

[45] “hg38 delly exclusion regions in bed format,” https://gist.github.com/chapmanb/4c40f961b3ac0a4a22fd, accessed: 2018-8-6.

[46] “Dwgsim,” https://github.com/nh13/DWGSIM, accessed: 2018-04-26.

[47] B. K. Stöcker, J. Köster, and S. Rahmann, “Simlord: simulation of long read data,” Bioinformatics, vol. 32, no. 17, pp. 2704–2706, 2016.

[48] J. M. Zook, D. Catoe, J. McDaniel, L. Vang, N. Spies, A. Sidow, Z. Weng, Y. Liu, C. E. Mason, N. Alexander et al., “Extensive sequencing of seven human genomes to characterize benchmark reference materials,” Scientific data, vol. 3, p. 160025, 2016.

[49] “Novoalign,” www.novocraft.com.

[50] D. Karolchik, A. S. Hinrichs, T. S. Furey, K. M. Roskin, C. W. Sugnet, D. Haussler, and W. J. Kent, “The ucsc table browser data retrieval tool,” Nucleic acids research, vol. 32, no. suppl 1, pp. D493–D496, 2004.

[51] V. Kuleshov, “Probabilistic single-individual haplotyping,” Bioinformatics, vol. 30, no. 17, pp. i379–385, Sep 2014.

[52] J. G. Cleary, R. Braithwaite, K. Gaastra, B. S. Hilbush, S. Inglis, S. A. Irvine, A. Jackson, R. Littin, S. Nohzadeh-Malakshah, M. Rathod et al., “Joint variant and de novo mutation identification on pedigrees from high-throughput sequencing data,” Journal of Computational Biology, vol. 21, no. 6, pp. 405–419, 2014.

[53] A. R. Quinlan and I. M. Hall, “Bedtools: a flexible suite of utilities for comparing genomic features,”Bioinformatics, vol. 26, no. 6, pp. 841–842, 2010.

[54] J. Köster, “Rust-bio: a fast and safe bioinformatics library,” Bioinformatics, vol. 32, no. 3, pp. 444–446, 2015.

[55] J. Köster and S. Rahmann, “Snakemake—a scalable bioinformatics workflow engine,” Bioinformatics, vol. 28, no. 19, pp. 2520–2522, 2012.

[56] G. Navarro and M. Raffinot, “A bit-parallel approach to suffix automata: Fast extended string matching,” in Annual Symposium on Combinatorial Pattern Matching. Springer, 1998, pp. 14–33.

