## Supplementary Material for "Longshot: accurate variant calling in diploid genomes using single-molecule long read sequencing"

### Supplementary methods and data for "Longshot: accurate variant calling in diploid genomes using single-molecule long read sequencing"

Peter Edge and Vikas Bansal

#### Simulating a diploid genome

In order to simulate Illumina and PacBio reads, a diploid genome was generated from the hg19 reference genome (chromosomes 1-22). For simplicity, only SNVs were simulated. Heterozygous SNVs were placed uniformly at random at a rate of 0.001 and homozygous SNVs were placed uniformly at random at a rate of 0.0005. The SNV alleles were selected according to the genotype priors used by Li et al (43). The phase for heterozygous genotypes was selected uniformly at random. Two fasta sequences for each chromosome were generated using "bcftools consensus" to insert the alleles at SNVs from each haplotype into the reference sequence.

#### Estimating coverage from aligned reads

For both Illumina and PacBio whole-genome read datasets, the median coverage for each dataset was measured by sampling 100,000 random positions from the genome using `bedtools random` (53) and counting the number of aligned reads covering each position (passing samtools flag filter `3844`). Then, the median value of the measured coverages was taken. For the simulated data, the median coverage was not measured, and instead the reported coverage is the read coverage that was generated for the simulation.

#### Identification of candidate SNVs

Longshot uses a simple (and standard) genotyping model for identifying candidate SNVs. For each position on the reference, the read bases piled up over that position are considered. The most frequent non-reference base is selected and denoted as the "1" allele. The three genotypes $G \in \{\text{"0/0"}, \text{"0/1"}, \text{"1/1"}\}$ are considered. Let $\mathbf{A}$ be the vector of allele observations over $\{\text{"0"}, \text{"1"}\}$. We wish to compute the probability of a non-reference genotype given the observed pileup alleles:

$$p(\text{"0/1"}, \text{"1/1"}|A) = 1 - p(\text{"0/0"}|\mathbf{A}) \; .$$

Using Bayes' rule,

$$p(G|\mathbf{A}) = \frac{p\left(\mathbf{A}|G)p(G)\right)}{\sum_G p(\mathbf{A}|G)p(G)} \tag{2}$$

The probability of the observed pileup alleles is the product of their respective probabilities:

$$p(\mathbf{A}|G) = \prod_{a \in \mathbf{A}} (a|G) \tag{3}$$

$$p(a|G) = \begin{cases} 1 - \varepsilon & \text{if } a = G_0 = G_1 \\ \varepsilon & \text{if } a \neq G_0 = G_1 \\ \frac{1}{2}(1-\varepsilon) + \frac{1}{2}\varepsilon & \text{otherwise} \end{cases} \tag{4}$$

$\varepsilon$ is the probability of a sequencing error to a specific base. Since quality values are of limited use for SMS reads, we use our own estimated base-mismatch emission scores for this value (see the section "alignment parameter estimation").

#### Finding non-repetitive anchors

For an SMS read overlapping a potential SNV site, non-repetitive "anchor sequences" to the left and right of the potential SNV site are identified to perform local realignment. For this, Longshot searches leftward (and rightward) from the SNV site to find a sequence of length $k$ (default value 6) in the reference sequence where the aligned SMS read matches the reference exactly. This implies that the k-mer in the SMS read was likely sequenced from the template without error and is aligned correctly. This assumption may not hold when the reference sequence is repetitive. For example, consider a 6-mer "AAAAAA" that matches between reference and SMS read perfectly. This may be a good anchor sequence if it is the only occurence of "AAAAAA" in the nearby reference sequence. It is a bad anchor if this is not the case; for instance, if it occurs inside an even larger homopolymer run of "A"s. We circumvent this issue by ignoring any potential anchor that occurs more than once on the reference sequence within the maximum anchor search window (100 bp by default). The rust-bio implementation of the BNDM algorithm is used to quickly perform this k-mer search(54)(56). If the leftward or rightward anchor search exceeds half the size of the maximum anchor search window, then that position on the reference and read is used as the anchor regardless of any other factors. This means that in the worst case with default parameters, a realignment window of 100 bp will be formed around the potential SNV. In order to avoid forming realignment windows at a locus with large gaps, if a insertion/deletion/refskip event of length $\geq 20$ is encountered near the window, the site is not realigned.

#### Pair-HMM realignment for clusters of SNVs

When multiple potential SNVs are located in close proximity, the realignment approach should consider the alternate haplotypes defined by these SNVs jointly rather than perform the local realignment for each SNV independently. This is especially important for false potential SNVs – it is common for a single true SNV to be misaligned in the original BAM so that the pileup-based scan identifies it as two or three potential heterozygous SNVs within a few bp of each other. Therefore, we use a simple approach to merge nearby SNVs into SNV "clusters". For every pair of adjacent potential SNVs, we merge them into the same SNV cluster (and merge their realignment windows) if their realignment window boundaries overlap. For a cluster with $n$ potential SNVs, we use the pair-HMM forward algorithm to realign against each of the $2^n$ possible short-haplotype sequences, which we will refer to as the set $\mathcal{H}$. For example, the possible haplotypes in the case of three potential SNVs are $\mathcal{H} = \{000, 001, 010, 100, 110, 101, 011, 111\}$ represented in bitstring form for the 3 SNV sites. We then use a Bayesian calculation similar to the single SNV case to calculate the probability of each possible short-haplotype $h \in \mathcal{H}$

$$p(h \mid read) = \frac{p(read \mid h)}{\sum_{h' \in \mathcal{H}} p(read \mid h')}$$

where $p(read \mid h)$ is calculated using the forward algorithm between the (multi-SNV) read-window and the "haplotype sequence" obtained by inserting into the reference sequence window each SNV in $h$. The short-haplotype $h_{max}$ maximizing $p(h \mid read)$ is selected and used to assign the call for each of the $n$ alleles. The quality value $q_i$ for each allele is calculated independently as:

$$q_i = \text{phred}\left(1 - \sum_{h_c} p(h_c \mid \text{read})\right)$$

where $h_c$ is the set of haplotypes in $\mathcal{H}$ for which $h_c[i] = h_{max}[i]$. In other words, it is 1 minus the sum of probabilities of all short haplotypes sharing the same best allele call in position $i$. The total computational complexity of all the realignments is $O(m^2 2^n)$, for read and haplotype windows of length $m$. For computational efficiency, we limit the cluster size ($n$) to a maximum value (default = 3) and break large clusters into smaller ones.

#### Priors on genotypes

Longshot uses the same approach as Li et al (43) to estimate the prior probability of a genotype $p(g)$. The prior probabilities for each SNV genotype (given the reference base) are derived assuming that heterozygous SNVs occur at a rate of 0.001 and homozygous SNVs occur at a rate of 0.0005. By default, Longshot differs from the Li et al approach in that a transition (Ts) mutation is assumed to occur at the same rate as a transversion (Tv) mutation. The prior probabilities can be specified as parameters to the software.

#### Haplotyping and measuring accuracy

Longshot produces a phased VCF as output, using the standard "phase set" (PS) notation to delineate phased haplotype blocks. To assess the accuracy of the haplotypes assembled by Longshot, the output VCF was first filtered to remove SNVs with a low "phase quality" ($PQ < 30$). This value is similar to the genotype quality (GQ), except that it represents the confidence in the most likely phased genotype ("0|0","0|1","1|0","1|1"). The GQ, on the other hand, combines the heterozygous phased genotypes together to represent the most likely unphased genotype ("0/0","0/1","1/1").

A switch error occurs when, at a single SNV, the phase (e.g. "0|1" or "1|0") of the assembled haplotype differs from the ground-truth haplotype with respect to the previously compared SNVs. A switch error indicates that the phase of the SNVs that follow will be different. If two consecutive switch errors occur such that the phase of only one SNV is incorrect, this corresponds to a "mismatch" error or a short switch error. We calculated the switch and mismatch error rate separately and report a single error rate by adding these two error rates.

To compare the phasing performance of short reads with long reads using Longshot, we used HapCUT2 to assemble haplotypes using short reads and variants called using Freebayes on the same set of reads (23). Variants identified from short read WGS can also be paired with long read data to assemble long haplotypes (13). This differs from Longshot, which requires no prior knowledge of SNVs (and genotypes) to assemble haplotypes. We compared the phasing accuracy of Longshot with this composite approach. For this, SNVs called using 30× Illumina WGS with the previously described filters were used with the extractHAIRS program to extract haplotype fragments in the PacBio reads mode (`--pacbio 1`). Then, the fragments were used to assemble haplotypes with HapCUT2 (`commit c2e66089bc8bb789e00db0b43f97bdf08e97c229`). The resulting haplotypes were filtered for a minimum mismatch quality (similar to the "phase quality" described for Longshot) of 30.

#### Separation of reads by haplotype

Let $H = (H_1, H_2)$ be the final pair of haplotypes output by Longshot. Assuming that the prior probability of a read originating from $H_1$ or $H_2$ is equal, the probability that a read $r$ was sampled from haplotype $H_1$ can be calculated as:

$$\frac{p(r|H_1)}{p(r|H_1) + p(r|H_2)}$$

The read is assigned to $H_1$ if this probability is at least $T$ (default value = 0.99), assigned to $H_2$ if the probability is $\leq 1 - T$ and left unassigned otherwise.

#### Alignment Parameter Estimation

In order to estimate allele probabilities using the pair-HMM, it is necessary to know the best alignment parameters for SMS reads. Specifically, the parameters are transition probabilities between every state in {MATCH, INSERTION, DELETION} (with outgoing probabilities for a state summing to 1) as well as emission probabilities for a pair of aligned bases. We use a single emission probability for matched bases, and a single emission probability for mismatched bases. In order to use parameters that accurately reflect the data, we estimate these probabilities directly from the alignments in the bam file. While the bam alignments are too inaccurate for sensitive genotyping, we can expect the alignments to roughly reflect the probabilities of insertion, deletion, and base mismatch errors for the reads. This approach also assumes that variants from the reference genome are significantly less common than read errors, which is true for SMS reads from the human genome. We perform a single scan over every CIGAR string and sequence in the BAM, and transition between MATCH, MISMATCH, and DELETION states according to the CIGAR string. The number of observed transitions from each state to itself or other states is counted and converted into probabilities by dividing by the total transitions out of the outgoing state. The aligned bases at each step in the CIGAR are tracked, and the total number of matching and mismatching bases are used to estimate the emission probability for matched vs. mismatched bases.

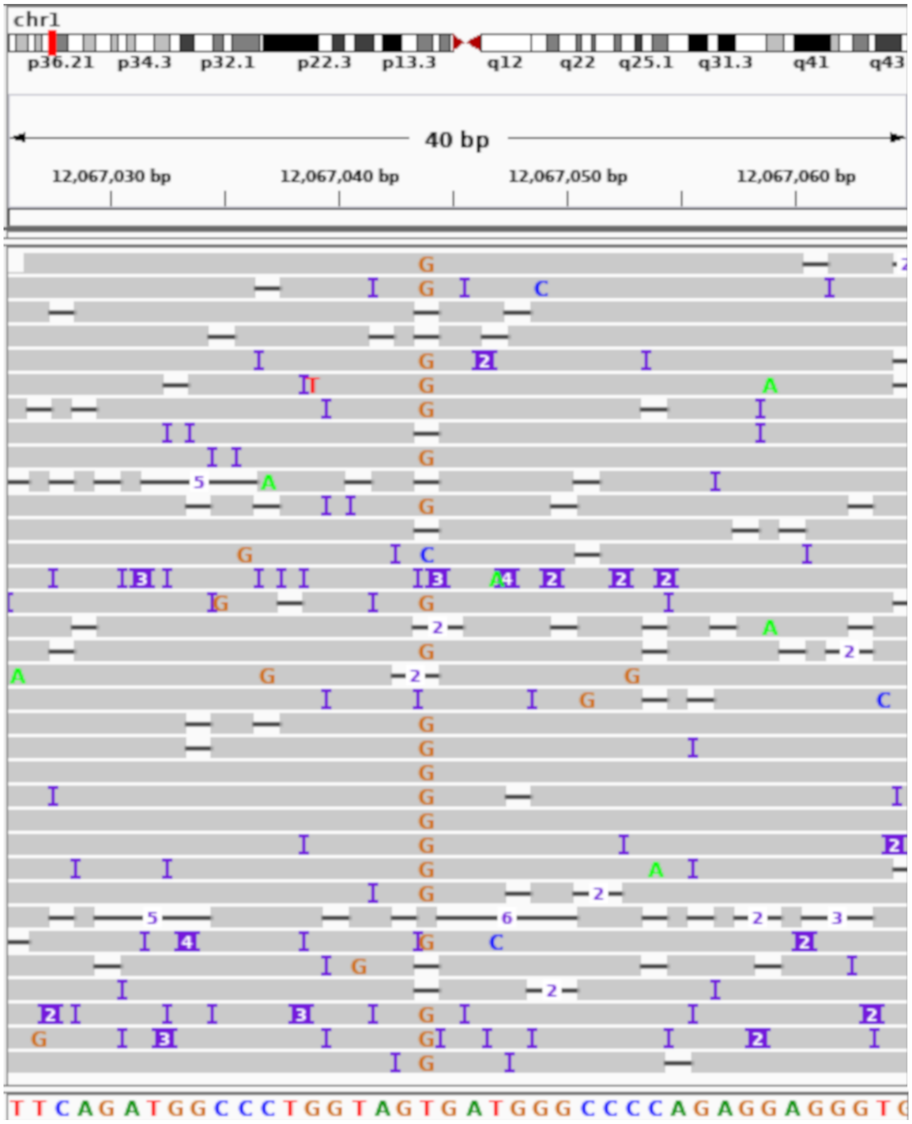


| Allele Call | Quality Score |
|---|---|
| G | 18.96 |
| G | 18.80 |
| G | 5.95 |
| G | 7.04 |
| G | 18.79 |
| G | 18.84 |
| G | 18.96 |
| G | 5.95 |
| G | 18.96 |
| G | 5.94 |
| G | 18.70 |
| G | 5.95 |
| G | 15.48 |
| G | 3.49 |
| G | 17.79 |
| G | 3.33 |
| G | 18.96 |
| G | 3.34 |
| T | 4.61 |
| G | 18.97 |
| G | 18.96 |
| G | 18.96 |
| G | 18.95 |
| G | 18.96 |
| G | 18.97 |
| G | 18.96 |
| G | 17.78 |
| T | 4.93 |
| G | 15.27 |
| G | 6.33 |
| G | 5.95 |
| G | 18.87 |
| G | 15.39 |
| G | 18.95 |

**Figure S1: Illustration of reference bias in SMS read alignments.** PacBio SMS reads covering a homozygous SNV (chr1:12,067,044 T→G) site in the individual NA12878 are shown (visualized using IGV). Frequent indel errors in the SMS reads, combined with a bias in the alignment algorithm to favor the reference allele, results in 3 reads that contain the reference allele at the SNV site, 1 read containing a C, and 9 reads that contain a deletion. As a result, the variant can incorrectly be called as a heterozygous SNV. Realigning each read to both the reference allele and the alternate allele and selecting the most likely alignment can ameliorate this bias. To the right of each read, the most likely allele call and quality value calculated by Longshot using the Pair-HMM realignment strategy is shown. All reads, except two reads with very low quality values ($< 5$), support the 'G' allele resulting in the correct homozygous SNV call.

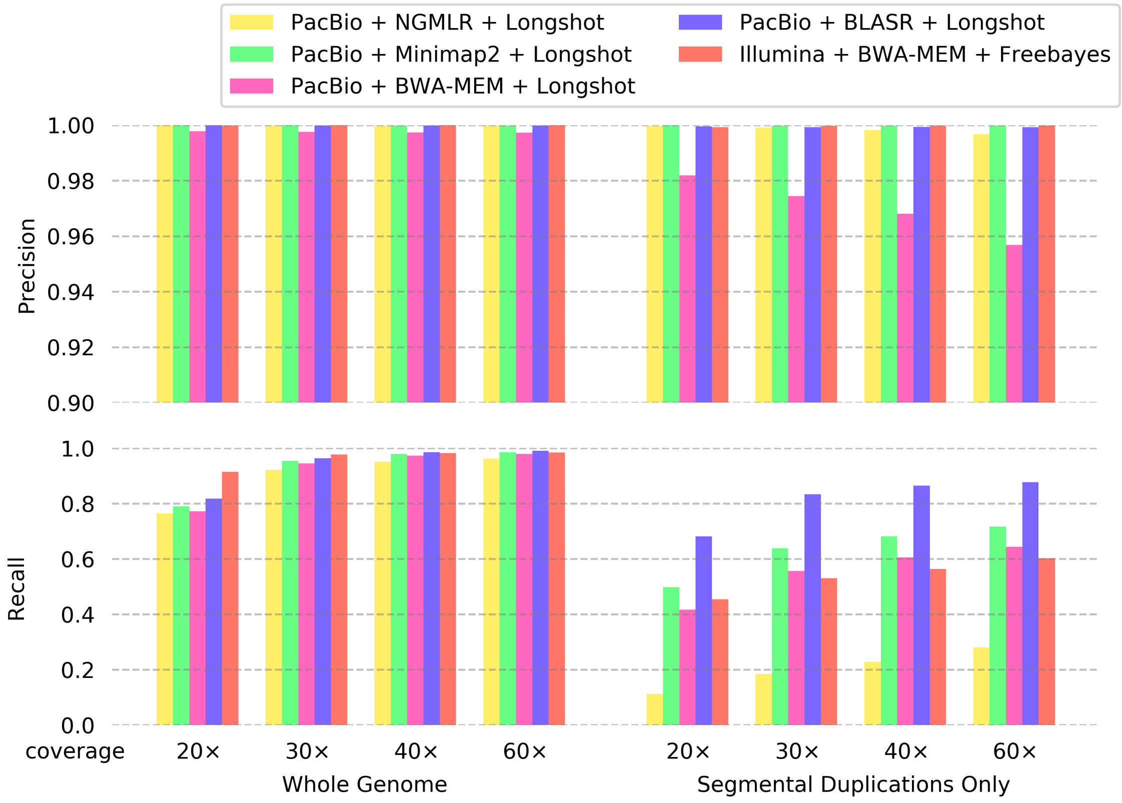


**Figure S2: Comparison of precision and recall of SNV calling using different long-read mapping tools.** Simulated Illumina and PacBio reads were generated at multiple coverages (20×, 30×, 40× and 60×) from chromosomes 1-22 (hs37d5 reference genome) and variants were called using either Longshot (PacBio) or Freebayes (Illumina). SMS reads were mapped with multiple mapping tools (NGMLR, BWA-MEM, MINIMAP2, and BLASR) for comparison. Precision **(top)** and Recall **(bottom)** of called variants were assessed across the entire chromosome **(left)** and within segmental duplications with 95% or greater sequence identity **(right)**.

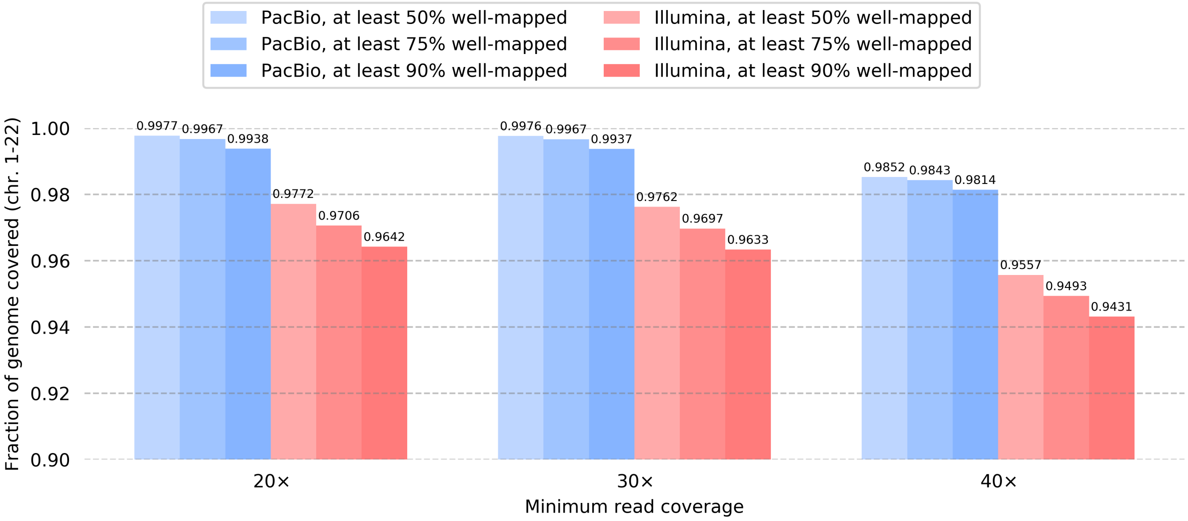


**Figure S3: Comparison of the mappability of short reads with long reads using simulated data.** Illumina short reads and PacBio SMS long reads were each simulated at 60× coverage and mapped to the genome with BWA-MEM and BLASR, respectively. For every position in the genome, the coverage of primary read mappings was assessed. The positions in the genome were filtered for those with at least 20×,30×, and 40× coverage of primary read mappings. Of those positions, it was determined what fraction of the mappings were “well-mapped”, or passing standard filters and having MAPQ $\geq$ 30. The number of positions meeting the minimum coverage cutoff and also meeting a minimum “well-mapped read” cutoff of at least 50%, 75%, 90% are shown as a fraction of total genomic positions (excluding “N” positions).

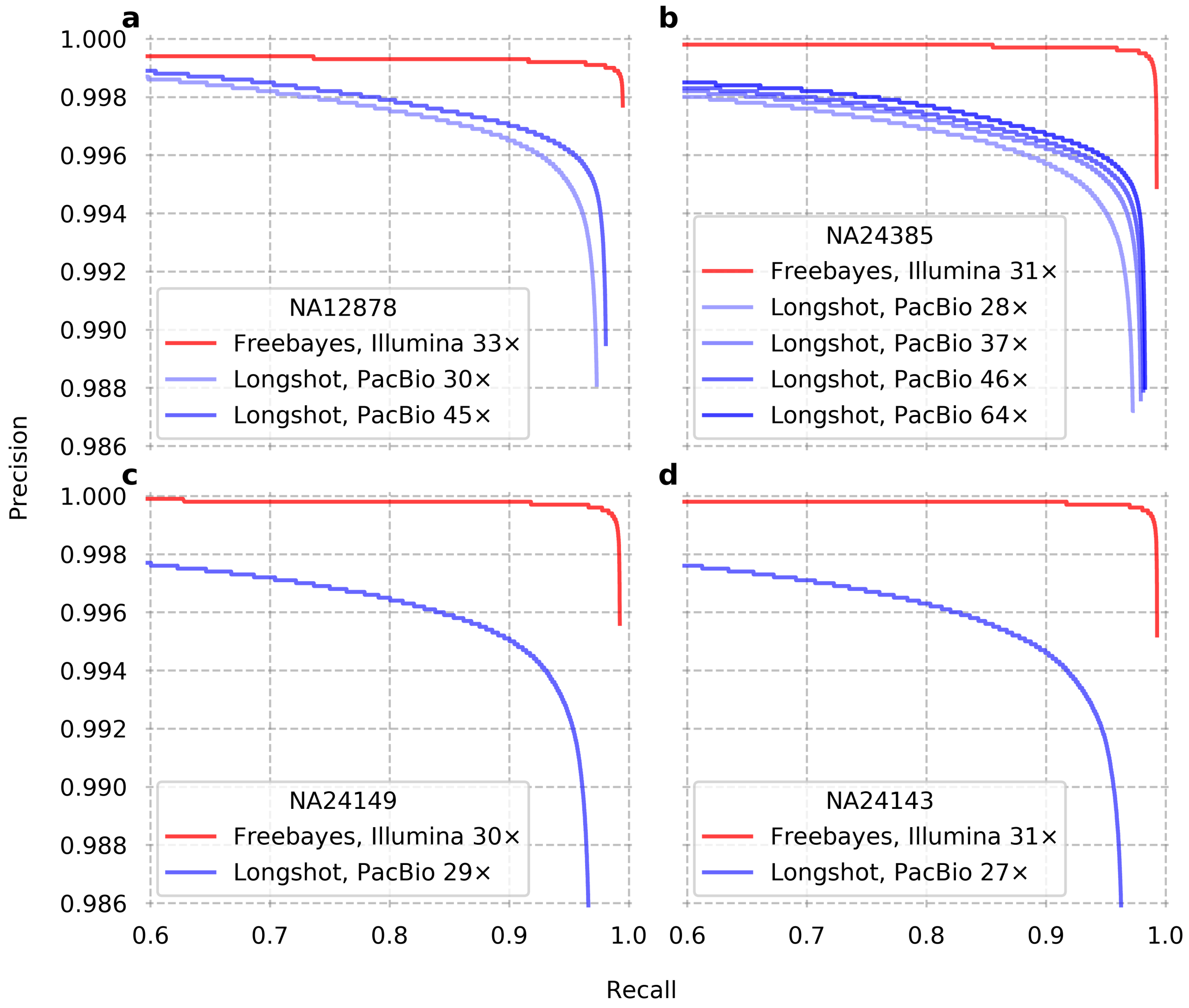


**Figure S4: Precision-Recall curve for SNV calling on four individuals: NA12878, NA24385, NA24149, and NA24143.** For each individual, variants were called from Illumina short reads using Freebayes and from whole-genome PacBio SMS reads using Longshot. Precision and recall were calculated using GIAB SNV calls within high-confidence regions. Points on each curve are obtained by varying the minimum Genotype Quality (GQ) cutoff.

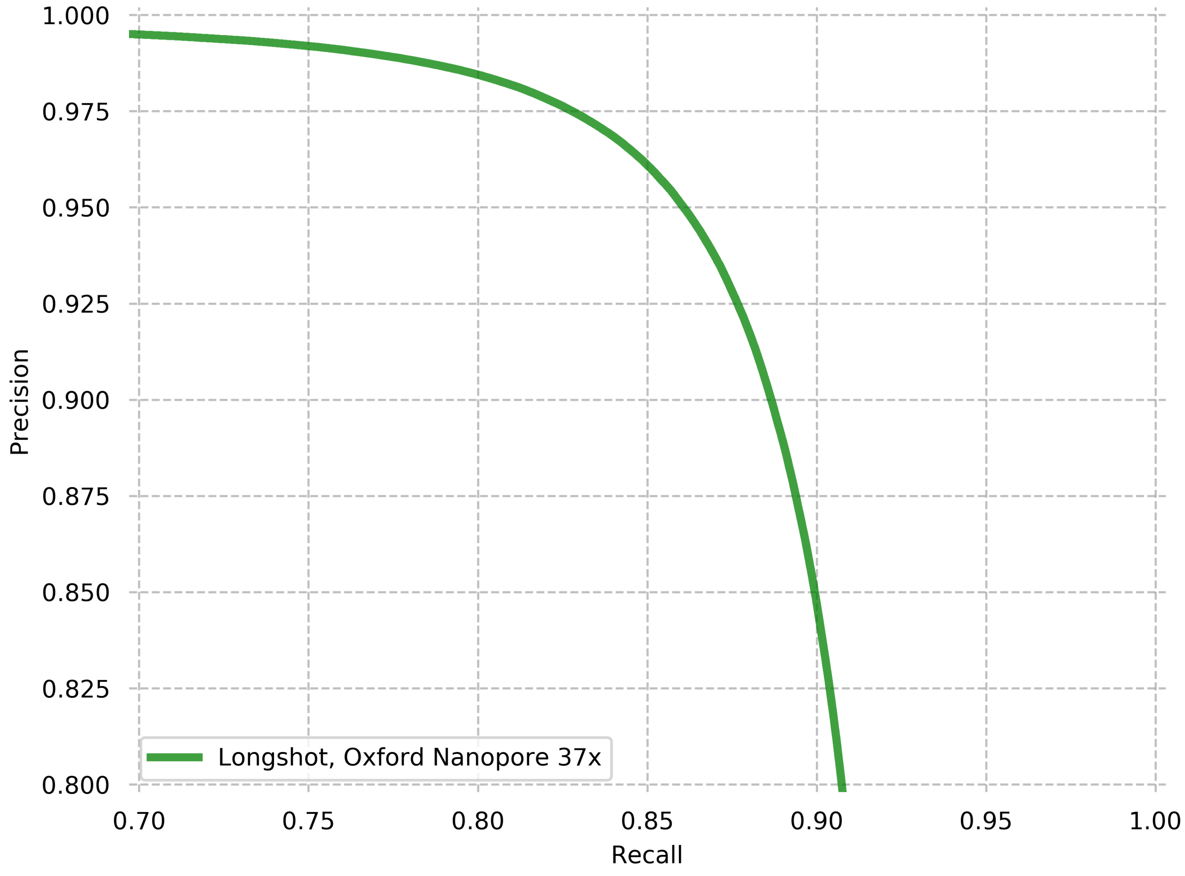


**Figure S5: Precision-Recall curve for SNV calling on NA12878 using 37× coverage Oxford Nanopore data.** Precision and recall were calculated using GIAB SNV calls within high-confidence regions. Points on the curve were obtained by varying the minimum Genotype Quality (GQ) cutoff.

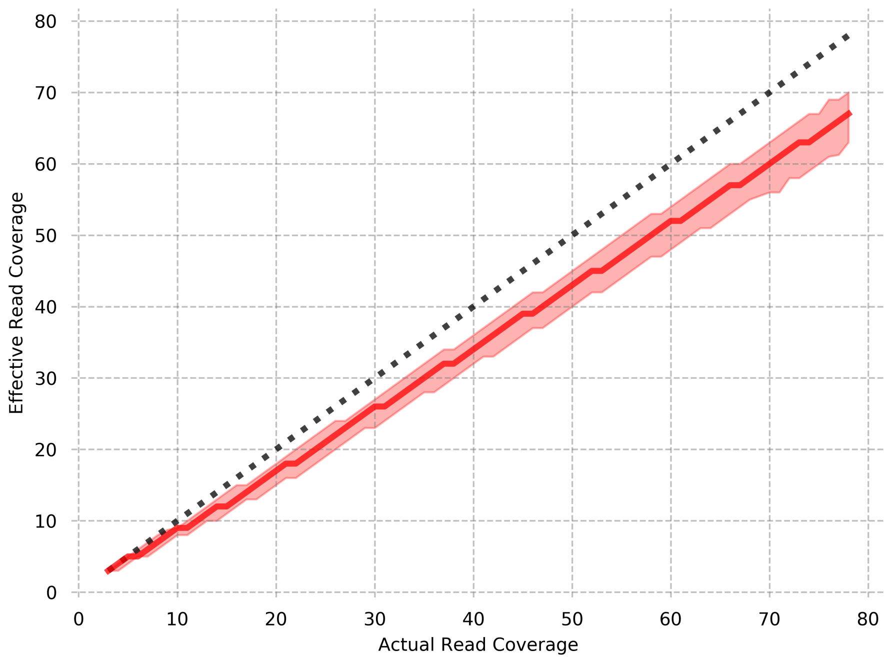


**Figure S6: Actual vs effective read coverage in PacBio SMS data**. At each SNV site, alleles for which the quality value estimated by the allelotyping method in Longshot is lower than a threshold (default = 7) are discarded; this reduces the effective read coverage. The data shown here is for SNV sites from the 45× coverage NA12878 PacBio dataset (chromosome 1 only). The median effective read coverage (as well as 1st and 3rd quartiles) is plotted for variant sites with the same actual read coverage.

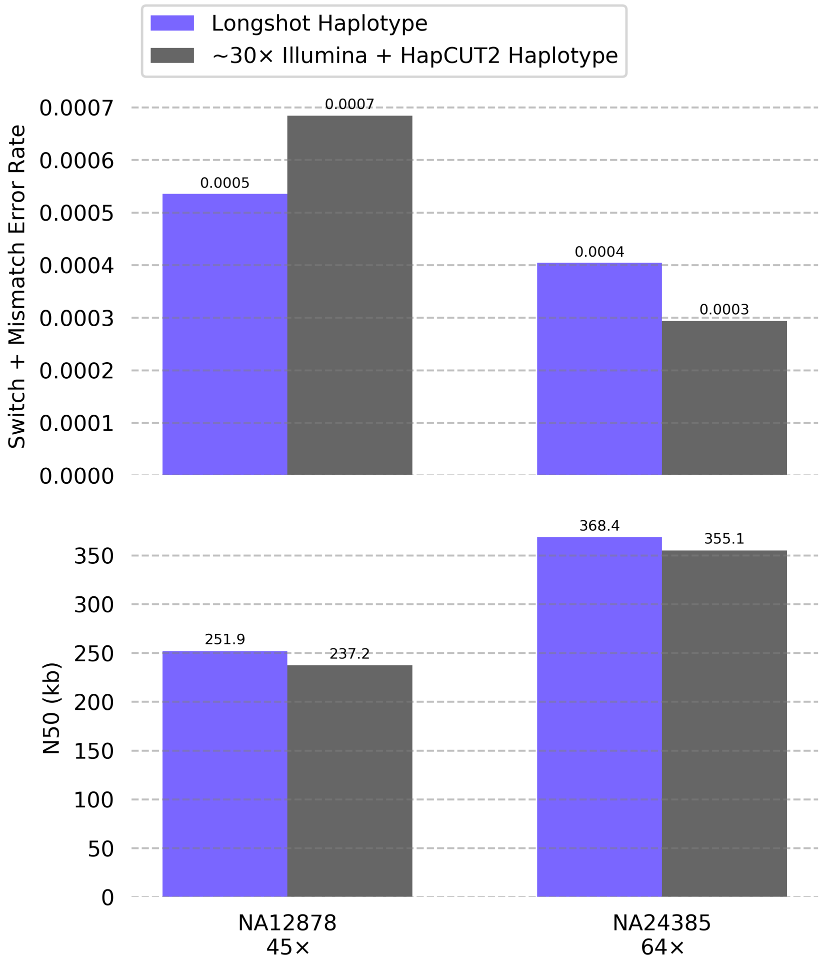


**Figure S7: Comparison of haplotypes assembled with Longshot and those assembled with HapCUT2.** Haplotypes were assembled for NA12878 (45× coverage SMS reads) and NA24385 (64× coverage SMS reads) using Longshot and HapCUT2. Longshot does not require prior knowledge of where SNVs, while for HapCUT2, variants identified from ∼30× coverage Illumina sequencing were used as input. The accuracy of the resulting haplotypes is measured using the combined switch error rate (long switches and mismatches), and the completeness is measured using the N50 of the haplotypes.

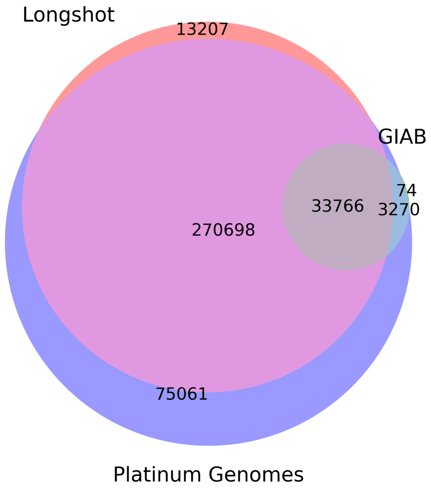


**Figure S8: Comparison of Platinum Genomes variant calls (outside GIAB confident regions) with Longshot variants for NA12878.** Variants were called on chr1-22 with Longshot using 45× coverage PacBio SMS reads for NA12878. The Longshot variants, GIAB variants, and Platinum Genomes variants were each filtered for regions that are outside GIAB confident regions, but inside the Platinum Genome called regions. 77% of SNVs in these difficult-to-call regions are shared between the PG calls and the Longshot calls.

| Genome | Read Coverage | SNVs called | Precision | Recall | Outside GIAB high-confidence | Run time (hours) |
|---|---|---|---|---|---|---|
| NA12878 | 30× | 3514112 | 0.994 | 0.958 | 513588 | 26:10 |
| NA12878 | 45× | 3564491 | 0.995 | 0.973 | 518698 | 35:33 |
| NA24385 (AJ son) | 28× | 3581651 | 0.993 | 0.960 | 659519 | 36:50 |
| NA24385 (AJ son) | 37× | 3625814 | 0.993 | 0.972 | 665091 | 40:44 |
| NA24385 (AJ son) | 46× | 3639281 | 0.993 | 0.976 | 664863 | 49:45 |
| NA24385 (AJ son) | 64× | 3648302 | 0.994 | 0.979 | 664291 | 68:51 |
| NA24149 (AJ father) | 29× | 3525248 | 0.992 | 0.951 | 774314 | 33:37 |
| NA24143 (AJ mother) | 27× | 3523724 | 0.992 | 0.944 | 770226 | 30:26 |

**Table S1: Summary of SNVs called using Longshot on whole-genome PacBio SMS data for multiple GIAB individuals.** Only variants called on the autosomes (chromosome 1-22) are reported. For the AJ son individual, variants were called at four different levels of coverage by downsampling. Precision and recall were calculated inside high-confidence GIAB regions using the GIAB variant set for each individual as the ground truth.

| Genome | False Positives (FP) | FP Enrichment | False Negatives (FN) | FN Enrichment |
|---|---|---|---|---|
| Misgenotyped SNV | 0.049 | - | 0.019 | - |
| Near Indel | 0.718 | 190.43 | 0.068 | 18.01 |
| In homopolymer | 0.317 | 5.51 | 0.205 | 3.56 |
| In homopolymer but not near indel | 0.025 | 0.44 | 0.165 | 2.93 |
| In STR | 0.008 | 26.96 | 0.004 | 11.95 |
| In LINE | 0.287 | 1.33 | 0.213 | 0.99 |
| In SINE | 0.124 | 0.80 | 0.222 | 1.44 |

**Table S2: Fractions of False Positive (FP) and False Negative (FN) variant calls that were misgenotyped or coincide with genomic features.** Variants were called using Longshot on chr1 using 45× coverage SMS reads for NA12878. The FP and FN variants were determined with respect to the ground truth (GIAB variant set within GIAB confident regions). In order to reason about why the FPs and FNs occurred, the fraction of those variants that fall into different categories was counted. For comparison, random positions from the GIAB confident regions were selected and subjected to the same analysis. The fold enrichment compared to the random positions is shown as a measure of the significance.

| Genome | Read Coverage | Precision (known indels not filtered) | Precision (known indels filtered out) | Recall (known indels not filtered) | Recall (known indels filtered out) |
|---|---|---|---|---|---|
| NA12878 | 45× | 0.995 | 0.997 | 0.973 | 0.968 |
| NA24385 (AJ son) | 64× | 0.994 | 0.996 | 0.979 | 0.973 |
| NA24149 (AJ father) | 29× | 0.993 | 0.995 | 0.947 | 0.942 |
| NA24143 (AJ mother) | 27× | 0.993 | 0.995 | 0.939 | 0.935 |

**Table S3: Improvement in variant precision by filtering out SNVs near known indel variants.** Most false positive (FP) variants called with Longshot occur at true indel variant sites that are mistaken as SNVs. Filtering out SNVs occuring within 5 bp of known indel sites (Mills + 1000G Indel variant set from the GATK bundle) results in improved precision at the cost of a small reduction in recall.

| Genome | Technology | Coverage | Method | Precision | Recall |
|---|---|---|---|---|---|
| NA12878 | PacBio | 45x | Longshot | 0.9948 | 0.9727 |
| NA12878 | PacBio | 45x | WhatsHap | 0.9738 | 0.9593 |
| NA12878 | PacBio | 45x | MarginPhase | 0.9497 | 0.9147 |
| NA12878 | PacBio | 40x | DeepVariant | 0.9819 | 0.9739 |
| NA12878 | PacBio | 45x | Clairvoyante | 0.9903 | 0.9705 |
| NA24385 | PacBio | 64x | Longshot | 0.9937 | 0.9792 |
| NA24385 | PacBio | 64x | Clairvoyante | 0.9894 | 0.9613 |
| NA12878 | ONT | 37x | LongShot | 0.9628 | 0.8478 |
| NA12878 | ONT | 37x | WhatsHap | 0.7131 | 0.7248 |
| NA12878 | ONT | 37x | MarginPhase | 0.8089 | 0.7686 |
| NA12878 | ONT | 37x | Clairvoyante | 0.9148 | 0.7518 |

**Table S4: Comparison of SNV calling accuracy across different methods on single-molecule WGS data for NA12878 and NA24385**. The precision and recall values for WhatsHap and MarginPhase were obtained from the bioRxiv preprint (http://dx.doi.org/10.1101/293944). Similarly, the precision and recall values for Clairvoyante were also obtained from the preprint (http://dx.doi.org/10.1101/310458, Table 7). For DeepVariant, the accuracy was obtained from Supplementary Data (Poplin et al., Nature Biotech., 2018).
